# Fluid flow-induced left-right asymmetric decay of *Dand5* mRNA in the mouse embryo requires Bicc1-Ccr4 RNA degradation complex

**DOI:** 10.1101/2020.02.02.931477

**Authors:** Katsura Minegishi, Benjamin Rothé, Kaoru R. Komatsu, Hiroki Ono, Yayoi Ikawa, Hiromi Nishimura, Emi Miyashita, Katsuyoshi Takaoka, Kana Bando, Hiroshi Kiyonari, Tadashi Yamamoto, Hirohide Saito, Daniel B. Constam, Hiroshi Hamada

## Abstract

Molecular left-right (L-R) asymmetry is established at the node of the mouse embryo as a result of the sensing of a leftward fluid flow by immotile cilia of perinodal crown cells and the consequent degradation of *Dand5* mRNA on the left side. We here examined how the fluid flow induces *Dand5* mRNA decay. We found that the 3’ untranslated region (3’-UTR) of *Dand5* mRNA is necessary and sufficient for the left-sided decay and is responsive to the flow direction, loss of the cation channel Pkd2, and Ca^2+^. The 200-nucleotide proximal-most portion of the 3’-UTR, which is conserved among mammals, is essential for the asymmetric mRNA decay and binds Bicc1, an RNA binding protein specifically expressed at the node. Bicc1 preferentially recognizes GAC and GACR sequences in RNA, and these motifs are enriched in the 200-nucleotide region of the *Dand5* 3’-UTR. The Cnot3 component of the Ccr4-Not deadenylase complex interacts with Bicc1 and is also required for *Dand5* mRNA decay at the node. Our results thus suggest that leftward fluid flow induces binding of Bicc1 to the 3’-UTR of *Dand5* mRNA in crown cells on the left side of the node, and that consequent recruitment of Ccr4-Not mediates mRNA degradation.

## INTRODUCTION

The first step in establishment of left-right (L-R) asymmetry in vertebrate embryos is known as the L-R symmetry breaking event. In fish, frog, and mouse embryos, the breaking of L-R symmetry requires motile cilia that generate a unidirectional fluid flow at the L-R organizer, which corresponds to Kupffer’s vesicle in zebrafish, the gastrocoel roof plate in *Xenopus*, and the ventral node in mouse ^1, 2^. Whereas this fluid flow, known as nodal flow in the mouse, is essential for L-R symmetry breaking, its mechanism of action remains unknown. It may transport an unknown chemical determinant or it may generate a mechanical stimulus, either of which would then be sensed by the embryo.

Genetic studies have indicated that nodal flow is sensed by the transient receptor potential (TRP) cation channel Pkd2 ^3^ and its partner protein Pkd1l1 ^4^ present in immotile cilia of crown cells at the periphery of the node ^5^. Flow sensation by Pkd2 is thought to result in the entry of Ca^2+^ and the consequent degradation through an unknown mechanism of the mRNA for Dand5 (also known as Cerl2 or Cer2) in crown cells on the left side of the node ^1, 6^. Given that Dand5 is an antagonist of the extracellular signaling molecule Nodal, which is required for correct L-R patterning ^7^, suppression of *Dand5* mRNA on the left side results in increased Nodal activity in crown cells on this side ^8^. Flow-induced down-regulation of *Dand5* mRNA specifically on the left side is the earliest molecular L-R asymmetry that has been detected at the node, and it in turn induces asymmetric expression of *Nodal* in the lateral plate ^8^.

The specific factors responsible for the suppression of *Dand5* mRNA have remained unidentified. The only RNA binding protein known to be required for correct L-R patterning is Bicc1 ^9^. Bicc1 binds specific mRNAs via a tandem repeat of three ribonucleoprotein K homology (KH) domains and thereby inhibits their translation in a manner that is facilitated by self-polymerization of a sterile α motif (SAM) at the COOH-terminus of the protein ^10, 11^. Studies in *Drosophila* have shown that Bicc1 also interacts with the Ccr4-Not complex—the major cytoplasmic deadenylase responsible for mRNA turnover in eukaryotes ^12, 13^—through direct binding to its Not3/5 subunit ^14^. In mammalian cells, lack of the Not3/5 ortholog Cnot3 is associated with increased mRNA stability ^15, 16^. However, a function for Bicc1 or the Ccr4-Not deadenylase complex in flow sensing, or in regulation of mRNA translation or decay, during L-R patterning has not been described.

We have now identified an element in the 3’ untranslated region (3’-UTR) of *Dand5* mRNA that is responsible for the asymmetric decay of this mRNA at the node of the mouse embryo in a manner dependent on Bicc1 and the Ccr4-Not subunit Cnot3.

## RESULTS

### A 3’-UTR Reporter Recapitulates the Asymmetric Distribution of *Dand5* mRNA at the Node

The L-R pattern of *Dand5* mRNA changes dynamically at the node of mouse embryos. At the early headfold stage, *Dand5* mRNA is induced specifically in crown cells on both sides of the node, and its distribution remains symmetric until the zero-somite stage (see Fig. 1). At the three-somite stage, the amount of *Dand5* mRNA has begun to decrease on the left side, resulting in a L-R asymmetric (R > L) distribution that is maintained at least until the six-somite stage. This decline in the abundance of *Dand5* mRNA on the left side coincides with a gradual increase in nodal flow ^8, 17^. Such observations have thus suggested that *Dand5* mRNA is degraded in crown cells on the left side of the node between the zero- and five-somite stages.

**Figure 1.**
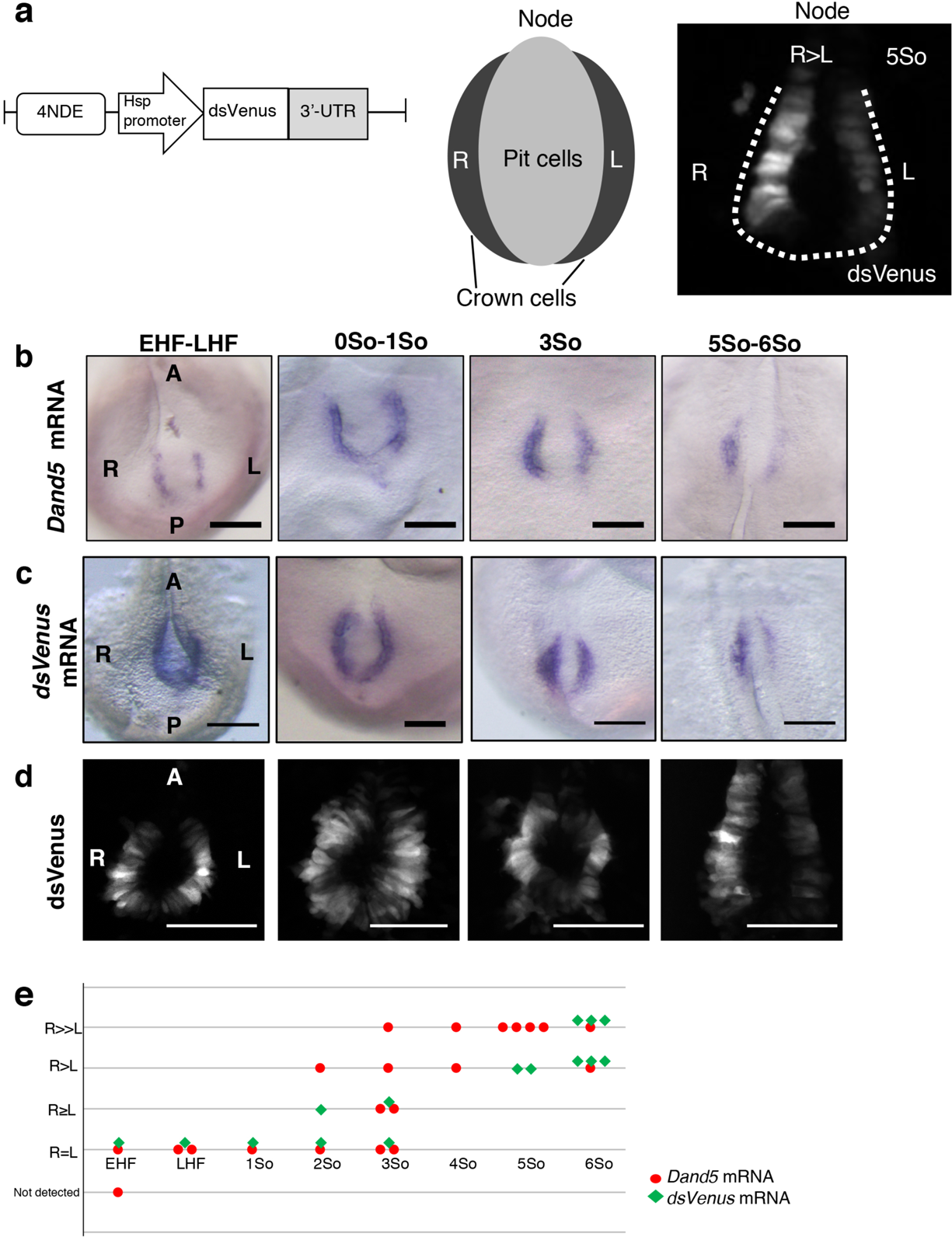
A 3’-UTR Reporter Recapitulates the Asymmetric Distribution of *Dand5* mRNA at the Node **a** Schematic representation of the *NDE-Hsp-dsVenus-3’-UTR* transgene (left) and of the relative localizations of pit cells and crown cells at the node of mouse embryos (middle). The transgene contains the ORF for dsVenus linked to the DNA sequence for the 3’-UTR (1.2 kb) of mouse *Dand5* mRNA, and its expression is driven by the mouse *Hsp68* promoter and four copies of the crown cell–specific enhancer (NDE) of mouse *Nodal*. A typical pattern of dsVenus fluorescence detected at the node of a transgenic embryo at the five-somite (5So) stage is also shown (right), with the dashed line indicating the node region. Note that the fluorescence in crown cells is highly L-R asymmetric (R > L). **b** Whole-mount in situ hybridization (WISH) analysis of *Dand5* mRNA at the node of wild-type mouse embryos at the indicated developmental stages. Note that *Dand5* mRNA in crown cells is bilaterally equal at the early headfold (EHF) to late headfold (LHF) stages as well as the zero- to one-somite stages, but is asymmetric (R > L) at the three-somite and five- to six-somite stages. A and P, anterior and posterior, respectively. Scale bars, 100 µm. **c** WISH analysis of *dsVenus* mRNA in *NDE-Hsp-dsVenus-3’-UTR* transgenic embryos at the indicated stages. Scale bars, 100 µm. **d** Fluorescence images of dsVenus at the node of transgenic embryos at the indicated stages. Scale bars, 100 µm. **e** Comparison of L-R asymmetry of *Dand5* mRNA in wild-type embryos and *dsVenus* mRNA in transgenic embryos at various developmental stages. Each point corresponds to one embryo. Note that the pattern of *dsVenus* mRNA recapitulates that of *Dand5* mRNA.

We first tested if the 3’-UTR of *Dand5* mRNA is able to recapitulate the distribution pattern of the mRNA at the node with the use of a transgene, *NDE-Hsp-dsVenus-3’-UTR*, that contains the open reading frame (ORF) for destabilized Venus (dsVenus) linked to the 1.2-kb DNA sequence corresponding to the 3’-UTR of *Dand5* mRNA (Fig. 1a). Expression of the *dsVenus-3’-UTR* cassette is driven by the mouse *Hsp68* promoter and the crown cell–specific enhancer (NDE) of mouse *Nodal*. As expected, *NDE-Hsp-dsVenus-3’-UTR* transgenic embryos expressed dsVenus specifically in node crown cells. Both *dsVenus* mRNA (Fig. 1c) and dsVenus fluorescence (Fig. 1a and 1d) were bilaterally equal at the early headfold and zero-somite stages but were asymmetric (R > L) at the three- and five-somite stages, recapitulating the pattern of *Dand5* mRNA (Fig. 1b and 1e). Hereafter, asymmetry of dsVenus fluorescence was examined at the five- to six-somite stages unless indicated otherwise.

### The 3’-UTR of *Dand5* mRNA Responds to the Direction of Fluid Flow in a Pkd2-Dependent Manner

To test whether the *NDE-Hsp-dsVenus-3’-UTR* transgene is regulated by nodal flow, we introduced it into *iv/iv* mutant embryos, which lack the fluid flow ^18^. In such embryos (n = 3/3), the transgene failed to give rise to an asymmetric (R > L) distribution of dsVenus fluorescence (Fig. 2a). Mouse embryos in culture develop L-R asymmetry in response to the direction of an artificial fluid flow ^19^. Exposure of wild-type (WT) embryos harboring the transgene to an artificial leftward fluid flow resulted in the development of R > L asymmetry of dsVenus fluorescence (n = 4/4) (Fig. 2b–2d). By contrast, such embryos exposed to an artificial flow directed toward the right side developed bilaterally symmetric (R = L) dsVenus fluorescence (n = 3/3). Exposure of *iv/+* or *iv/iv* embryos to the artificial rightward flow resulted in the development of R < L (n =^20^ 3/6), R = L (n = 2/6), or R > L (n = 1/6) patterns of dsVenus fluorescence in the former embryos and in R < L (n = 4/7), R = L (n = 1/7), and R > L (n = 2/7) patterns in the latter (Fig. 2b–2d). These results collectively suggested that the 3’-UTR of *Dand5* mRNA responds to the direction of fluid flow.

**Figure 2.**
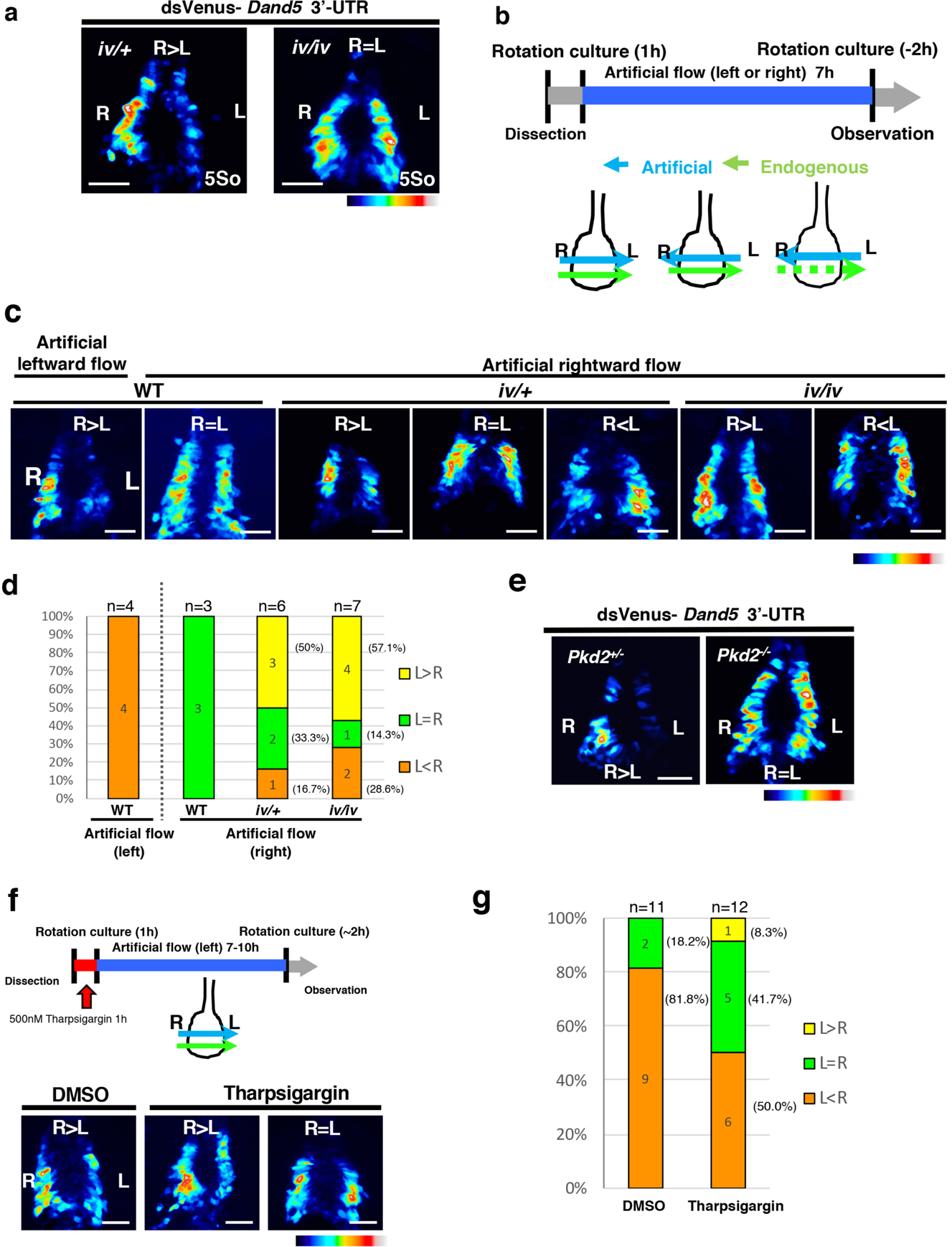
The 3’-UTR of *Dand5* mRNA Responds to the Direction of Fluid Flow in a Pkd2-Dependent Manner **a** Fluorescence of dsVenus at the node of *Dnah11*^iv/+^ or *Dnah11*^iv/iv^ embryos harboring the *NDE-Hsp-dsVenus-3’-UTR* transgene at the five-somite stage. The color scale indicates the intensity of dsVenus fluorescence, with white and black corresponding to the highest and lowest levels, respectively. Scale bars, 50 µm. **b** Protocol for artificial flow experiments. *Dnah11^+/+^* (WT), *Dnah11*^iv/+^, or *Dnah11*^iv/iv^ embryos harboring the *NDE-Hsp-dsVenus-3’-UTR* transgene were cultured under the influence of a rightward or leftward artificial flow from the <u>late</u> headfold to five-somite stages. **c** Fluorescence of dsVenus at the node of embryos of the indicated genotypes after exposure to the leftward or rightward artificial flow as in (**b**). Scale bars, 50 µm. **d** Summary of the L-R patterns of dsVenus fluorescence at the node of embryos as in (**c**). The numbers of embryos showing each pattern are indicated within the bars. **e** Fluorescence of dsVenus at the node of *Pkd2*^+/−^ or *Pkd2*^−/−^ embryos harboring the *NDE-Hsp-dsVenus-3’-UTR* transgene at the five-somite stage. Scale bars, 50 µm. **f** Determination of the effect of thapsigargin (500 nM) or dimethyl sulfoxide (DMSO, vehicle) treatment before imposition of a leftward artificial flow on dsVenus fluorescence at the node of WT embryos harboring the *NDE-Hsp-dsVenus-3’-UTR* transgene. Scale bars, 50 µm. **g** Summary of the effect of tharpsigargin on the L-R pattern of dsVenus fluorescence as in (**f**). The numbers of embryos showing each pattern are indicated within the bars.

*Pkd2* encodes an ion channel that allows the passage of cations including Ca^2+ 20^ and is required for L-R patterning in the mouse embryo ^3^. In *Pkd2*^−/−^ embryos (n = 3/3), the *NDE-Hsp-dsVenus-3’-UTR* transgene was expressed bilaterally at the node (Fig. 2e). Similarly, treatment of WT embryos harboring the transgene for 1h with thapsigargin, which blocks Ca^2+^ release from the endoplasmic reticulum ^21^, resulted in half of the tested embryos (n = 6/12) failing to down-regulate the reporter mRNA on the left side in response to an artificial leftward flow, giving rise to L = R (n = 5/12) or R < L (n = 1/12) patterns of dsVenus fluorescence (Fig. 2f and 2g). Longer treatment with thapsigargin resulted in developmental arrest. Together, these results implicate Ca^2+^ in the flow-induced decay of *Dand5* mRNA.

### A Conserved 200-Nucleotide Region of the Proximal 3’-UTR Is Required for L-R Asymmetry of *Dand5* mRNA

To map the sequence within the 3’-UTR required for generation of L-R asymmetry of *Dand5* mRNA, we first compared the DNA sequences corresponding to the 3’-UTR (∼1.2 kb) among mammalian *Dand5* genes. We found that the 5’-most 200 nucleotide region was substantially conserved (Fig. 3a). To test whether this region is required for L-R asymmetric gene expression, we first deleted it from the *NDE-Hsp-dsVenus-3’-UTR* transgene (Fig. 3b). The resulting *NDE-Hsp-dsVenus-3’-UTRΔ200* transgene yielded bilaterally symmetric dsVenus fluorescence at the node (Fig. 3c). Promted by this result, we applied CRISPR-based editing to delete this 200 nucleotide region of the *Dand5* 3’-UTR in the mouse genome, thereby generating the *Dand5*^Δ200^ allele (Supplementary Fig. 1). WISH analysis revealed that whereas bilateral *Nodal* expression at the node was maintained, left-sided *Nodal* expression in lateral plate mesoderm (LPM) was lost in *Dand5*^Δ200/Δ200^ embryos (n = 3/3), as expected if *Dand5* mRNA remains stable on both sides of the node. Among eight *Dand5*^Δ200/+^ embryos examined, one embryo lost *Nodal* expression in LPM, suggesting that the *Dand5*^Δ200^ allele may act dominantly in rare instances. WISH analysis also showed that *Dand5* mRNA was not distributed asymmetrically in *Dand5*^Δ200/Δ200^ embryos (Fig. 3e).

**Figure 3.**
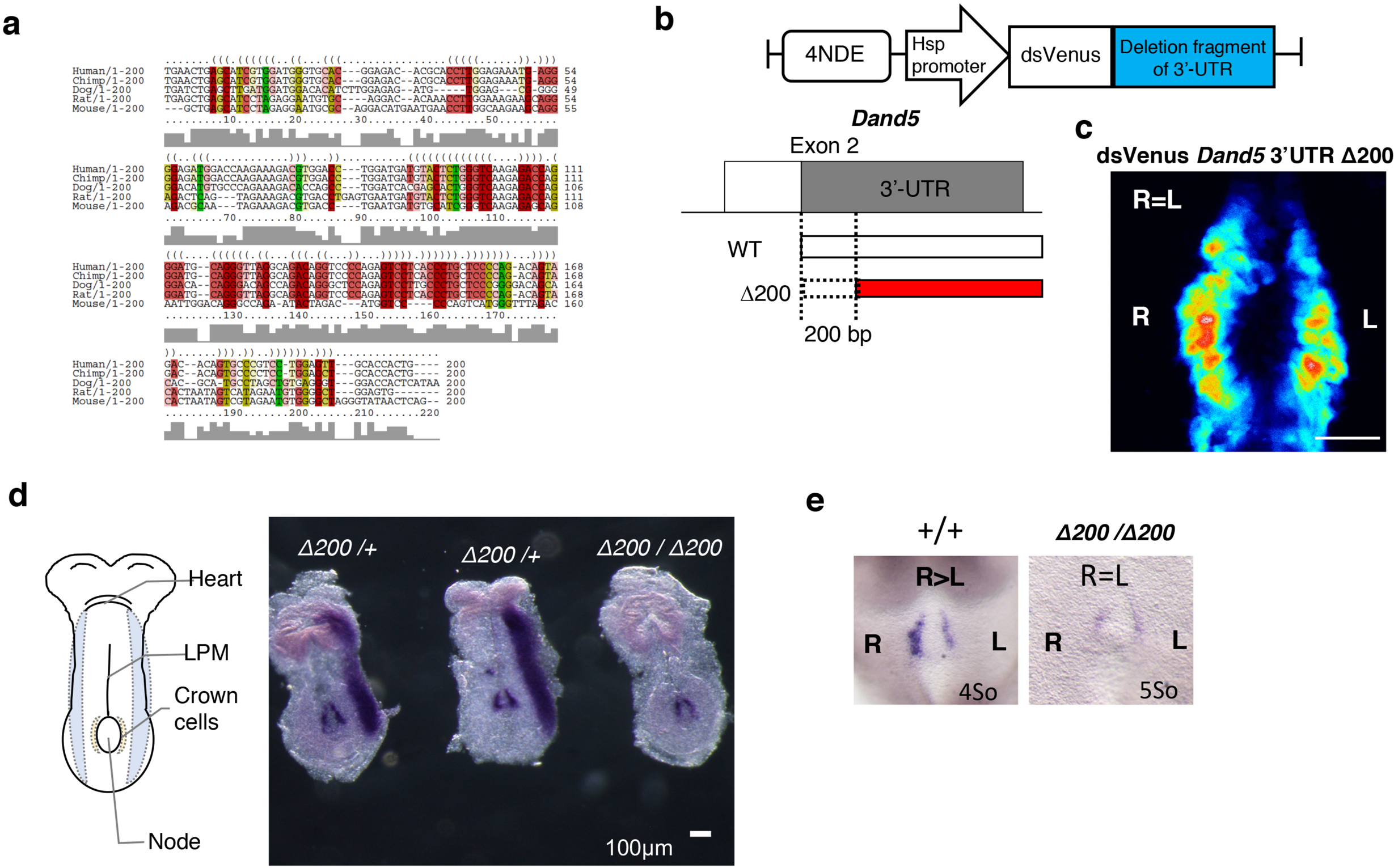
A 200-Nucleotide Conserved Proximal Region of the 3’-UTR Is Required for L-R Asymmetry of *Dand5* mRNA **a** Comparison of the DNA sequences corresponding to the 200-nucleotide most-proximal region of the 3’-UTR of *Dand5* mRNA among mammals. **b** Schematic representation of the *NDE-Hsp-dsVenus-3’-UTRΔ200* transgene, which lacks the 200-bp region corresponding to the mouse *Dand5* 3’-UTR in (a). **c** Fluorescence of dsVenus at the node of a mouse embryo harboring the *NDE-Hsp-dsVenus-3’-UTRΔ200* transgene at the five-somite stage. Scale bar, 50 µm. **d** WISH analysis of *Nodal* expression in *Dand5*^Δ200/+^ and *Dand5*^Δ200/Δ200^ embryos at embryonic day (E) 8.0. Left-sided expression of *Nodal* in LPM was lost in the *Dand5*^Δ200/Δ200^ embryo. Scale bar, 100 µm. **e** WISH analysis of *Dand5* expression at the node of *Dand5*^+/+^ and *Dand5*^Δ200/Δ200^ embryos at E8.0 the four- or five-somite stage. L-R asymmetry of *Dand5* mRNA at the node was lost in the *Dand5*^Δ200/Δ200^ embryo. See also Figure S1.

Together, these results indicated that the 200 nucleotide proximal region of the 3’-UTR is essential for the L-R asymmetric pattern of *Dand5* mRNA in node crown cells.

### Flow-Induced Decay of *Dand5* mRNA Is Dependent on Bicc1

We hypothesized that a specific factor (or factors) that is expressed in crown cells and binds to the proximal portion of the 3’-UTR mediates the decay of *Dand5* mRNA preferentially on the left side of the node. Among candidates for such a factor, we focused on Bicc1, an RNA binding protein with KH domains for RNA binding and a SAM domain for polymerization ^11, 22^, given that it is specifically expressed at the node of mouse embryos and that *Bicc1*^−/−^ embryos manifest laterality defects ^9^. Whole-mount immunofluorescence analysis revealed that pit cells and crown cells at the node express Bicc1 in an apparently L-R symmetric manner (Fig. 4a–4c). Moreover, whereas *Dand5* mRNA is localized predominantly at the apical side of crown cells ^23^, Bicc1 was detected both apically and basally (Fig. 4d). To evaluate potential genetic interaction between *Bicc1* and *Dand5*, we deleted the first exon of *Bicc1* with the CRISPR/Cas9 system (Supplementary Fig. Sa). The fluorescence of dsVenus in the resulting *Bicc1*^Δex1/Δex1^ embryos harboring the *NDE-Hsp-dsVenus-3’-UTR* transgene was bilaterally symmetric (Fig. 4f), suggesting that the decay of *Dand5* mRNA on the left side is impaired in the absence of Bicc1. Similar to our previous observations with *Bicc1*^−/−^ embryos ^9^, the basal body of motile cilia at the node failed to shift posteriorly in half of the *Bicc1*^Δex1/Δex1^ embryos examined (n = 3/6) (Supplementary Fig. 2b and 2c).

**Figure 4.**
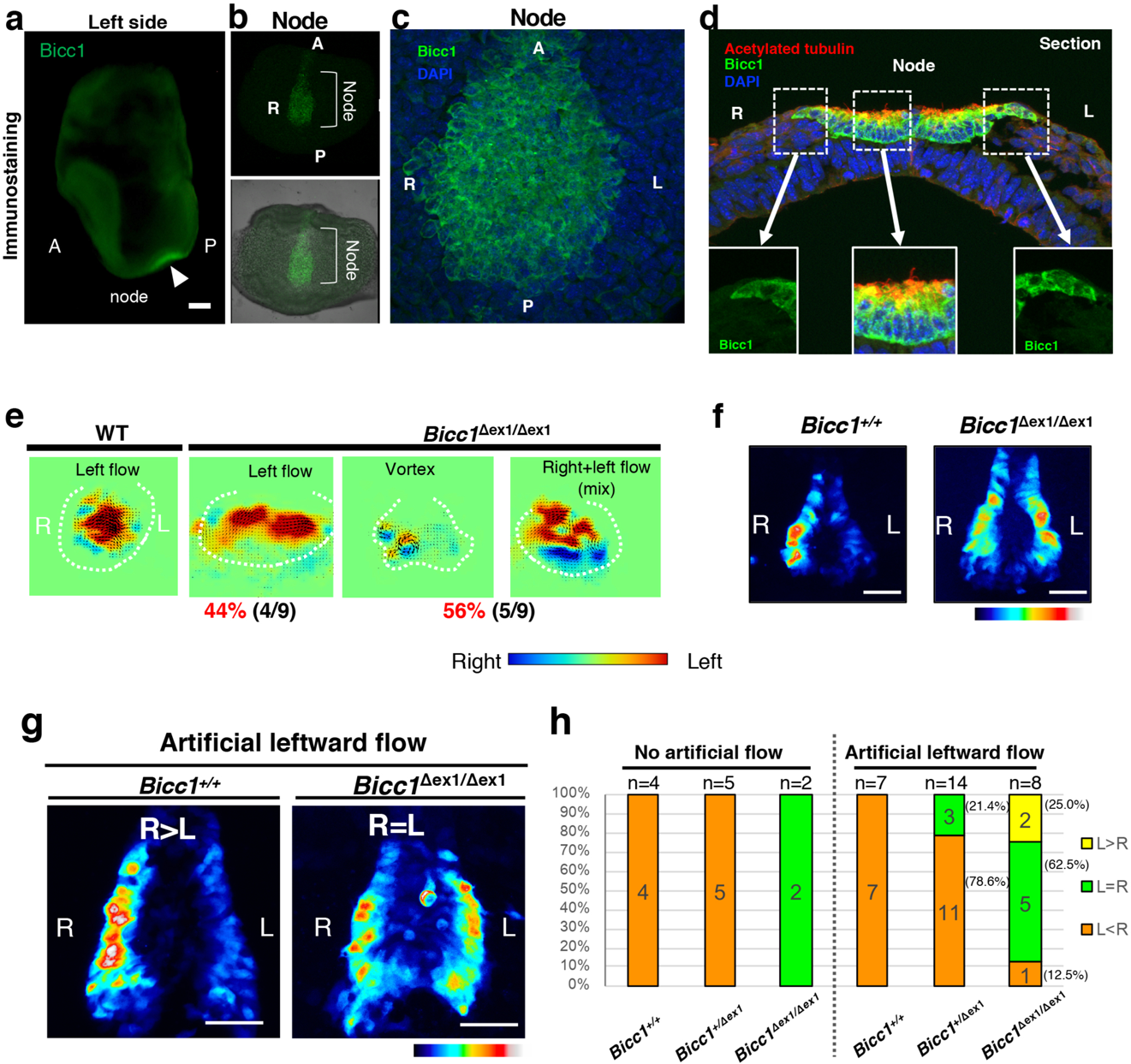
Flow-Induced Decay of *Dand5* mRNA Is Dependent on Bicc1 (**a**–**d**) Immunofluorescence staining of Bicc1 in node cells of the WT mouse embryo. Staining is shown for the whole embryo at E8.0 (a), for the node region at higher magnification (b and c), and for a transverse section of the node (d). Nuclei were stained with 4’,6-diamidino-2-phenylindole (DAPI) in (c) and (d), and acetylated tubulin was also immunostained in (d) to reveal cilia. Scale bar, 100 µm. **e** Nodal flow in WT and *Bicc1*^Δex1/Δex1^ mouse embryos at three- to five-somite stages as revealed by particle image velocimetry (PIV) analysis. The dashed white line indicates the outline of the node, small arrows in the node region indicate the direction and velocity of flow at those positions, and the relative color scale indicates the magnitude of flow velocity (leftward in red, rightward in blue). Note that leftward laminar flow was lost in about half of *Bicc1*^Δex1/Δex1^ embryos (n = 5/9), whereas the remaining mutant embryos (n = 4/9) showed normal leftward flow. **f** The L-R pattern of dsVenus fluorescence at the node of WT and *Bicc1*^Δex1/Δex1^ embryos harboring the *NDE-Hsp-dsVenus-3’-UTR* transgene at the X-somite stage. Scale bars, 50µm. **g** *Bicc1^+/+^* and *Bicc1*^Δex1/Δex1^ embryos harboring the *NDE-Hsp-dsVenus-3’-UTR* transgene were cultured in the presence of an artificial leftward flow as in Figure 2B, after which the L-R pattern of dsVenus fluorescence at the node was examined. Scale bars, 50 µm. **h** Summary of the L-R pattern of dsVenus fluorescence at the node of embryos of the indicated genotypes harboring the *NDE-Hsp-dsVenus-3’-UTR* transgene after culture under conditions of no or leftward artificial flow as in (g). Note that the artificial leftward fluid flow failed to rescue the impaired L-R pattern of dsVenus fluorescence in most of the *Bicc1*^Δex1/Δex1^ embryos (n = 7/8). See also Figure S2.

Consistent with this finding, the leftward flow at the node was disrupted in half of *Bicc1*^Δex1/Δex1^ embryos (n = 5/9), with the other half (n = 4/9 embryos) showing normal leftward flow (Fig. 4e). Nonetheless, an artificial leftward fluid flow failed to rescue the pattern of dsVenus fluorescence in seven of the eight *Bicc1*^Δex1/Δex1^ embryos examined, suggesting that the mutant embryos are unable to respond to the fluid flow (Fig. 4g and 4h).

### Bicc1 Binds to the 3’-UTR of *Dand5* mRNA

To assess whether the 200-nucleotide proximal portion of the 3’-UTR of *Dand5* mRNA mediates the binding of Bicc1, we forcibly expressed *dsVenus-3’-UTR* or *dsVenus-3’-UTRΔ200* constructs (similar to the corresponding transgenes but driven by a ubiquitous promoter) together with hemagglutinin epitope (HA)-tagged mouse Bicc1 in HEK293T cells. Immunoprecipitation of cell extracts with antibodies to HA followed by reverse transcription (RT) and quantitative polymerase chain reaction (qPCR) analysis of the immunoprecipitates and input extracts revealed that the *dsVenus-3’-UTR* mRNA was enriched by a factor of ∼8 in the HA-Bicc1 precipitates, whereas the *dsVenus-3’-UTRΔ200* mRNA or β-actin mRNA (negative control) showed no such enrichment (Fig. 5a–5d). Similar results were obtained by immunoprecipitation of endogenous Bicc1 from IMCD3 cells stably expressing the WT or mutant *dsVenus-3’-UTR* constructs (Supplementary Fig. 3a). Furthermore, a truncated form of Bicc1 lacking all KH domains (ΔKH) ^9^ failed to recruit the WT *dsVenus-3’-UTR* mRNA (Fig. 5d). In addition to the three tandem canonical KH domains, Bicc1 contains two interspersed KH-like (KHL) domains that lack the conserved GXXG motif (where at least one X is a basic residue) that is able to bind single-stranded RNA sequences of 4 nucleotides and orient them toward an adjacent groove for nucleobase recognition (Fig. 5e) (Nakel et al., 2010; Rothé et al., 2015; Nicastro et al., 2015). To evaluate the contribution of individual KH domains to RNA binding, we mutated the GXXG motif of each KH domain to GDDG ^24^ (Fig. 5f). Mutation of the KH2 domain alone impaired the binding of Bicc1 to the *dsVenus-3’-UTR* reporter mRNA to the same extent as did that of all three KH domains. By contrast, mutation of the KH1 or KH3 domain alone had no significant effect on binding, suggesting that the interaction relies largely on the KH2 domain. These results are consistent with the recent finding that the KH2 domain of *Xenopus* Bicc1 plays the major role in binding to *Cripto1* mRNA ^25^. Unexpectedly, however, despite efficient binding of HA-Bicc1 to the mRNA, the amount of dsVenus protein in HEK293T cells expressing the *dsVenus-3’-UTR* construct was unaffected by coexpression of HA-Bicc1 (Supplementary Fig. 3b). Together, these results suggest that binding of Bicc1 to the 3’-UTR of *Dand5* mRNA is mediated by the 200 nucleotide proximal portion of the 3’-UTR and the KH2 domain of Bicc1.

**Figure 5.**
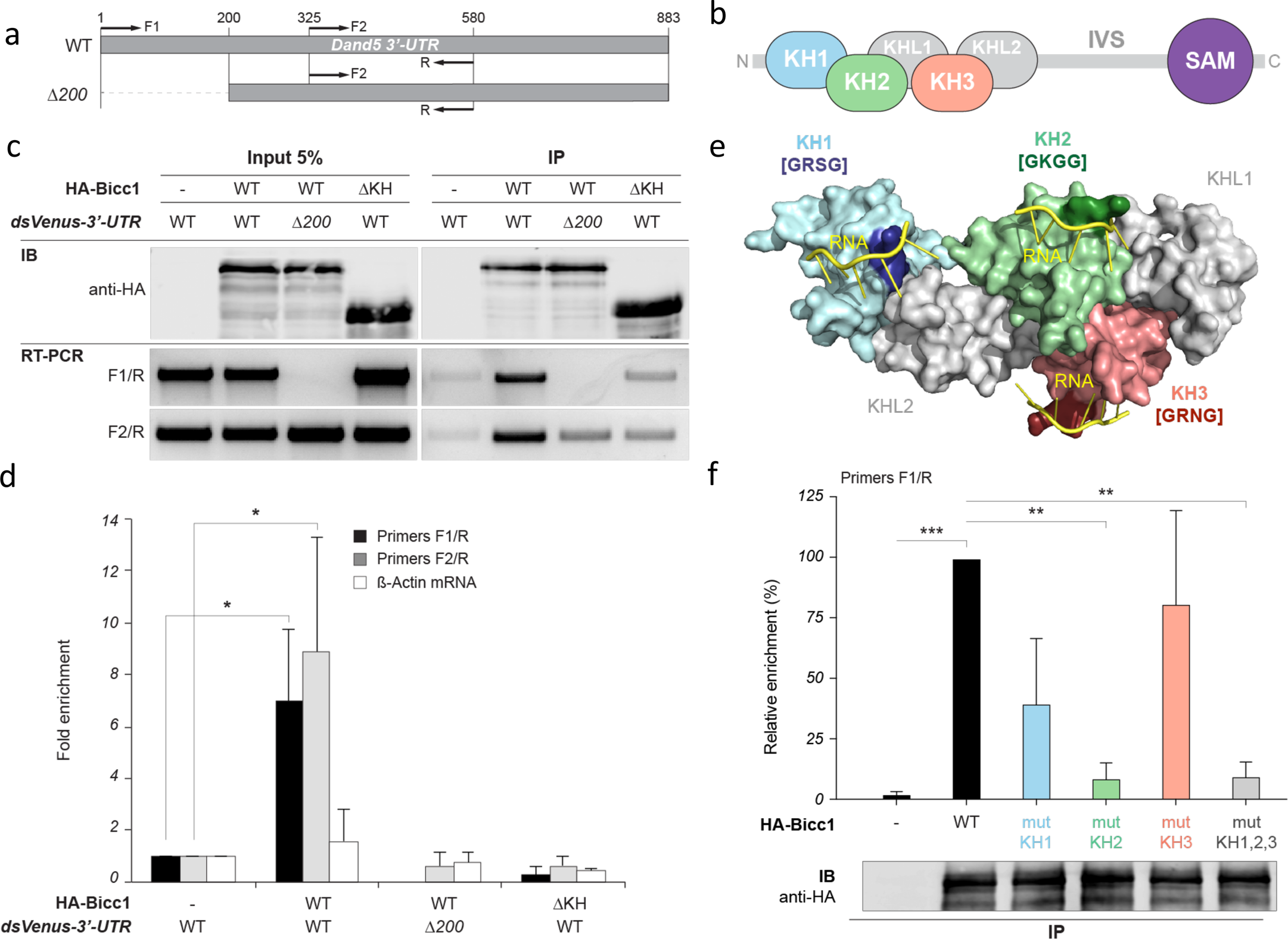
Bicc1 Binds to the 3’-UTR of *Dand5* mRNA via Its KH2 Domain **a** Annealing sites of PCR primers used to detect *Dand5* reporter (*dsVenus-3’-UTR* or *dsVenus-3’-UTRΔ200*) mRNAs by RT-PCR or -qPCR analysis. The primer pair F1/R detects only the WT construct, whereas the F2/R pair detects both WT and *Δ200* constructs. **b** Schematic of the domain organization of Bicc1. KH: K homology; KHL: KH-like; IVS: Intervening sequence; SAM: Sterile α motif. **c** Co-immunoprecipitation of mRNAs with HA-Bicc1 in HEK293T cells. Cytoplasmic extracts of cells expressing either of the reporter constructs in (a) as well as HA-Bicc1 (WT) or HA-Bicc1ΔKH were subjected to immunoprecipitation with antibodies to HA, and the resulting immunoprecipitates (IP) as well as portions of the input extracts were subjected to immunoblot (IB) analysis with antibodies to HA and to RT-PCR analysis with the PCR primers shown in (a). **d** RT-qPCR analysis of immunoprecipitates from experiments as in (b), but including β-actin mRNA as a negative control. Data represent the ratio of co-immunoprecipitated mRNA to the input and are expressed relative to the corresponding condition without HA-Bicc1. **e** Three-dimensional model of the region of mouse Bicc1 containing the three KH domains in surface representation. In each of the KH domains, putative RNA-binding GXXG motifs (where at least one X is a basic residue) are highlighted. **f** Immunoblot and RT-qPCR analyses as in (b) and (c) of immunoprecipitates prepared from HEK293T cells expressing the *dsVenus-3’-UTR* construct and wild-type HA-Bicc1 (WT) or mutant versions in which the GXXG motif of individual KH domains (mutKH1, mutKH2, or mutKH3) or all three KH domains (mutKH1,2,3) were replaced with GDDG. RT-qPCR data represent the ratio of co-immunoprecipitated mRNA to the input and are expressed as a percentage of the value for HA-Bicc1(WT). Data in (c) and (e) are means + SD from three independent experiments. *p < 0.05, **p < 0.01, ***p < 0.005 (Student’s t-test). See also Figures S3 and S4.

We next attempted to identify the RNA binding motifs for Bicc1 in order to further understand the relation between Bicc1 and the 3’-UTR of *Dand5* mRNA. We thus performed RNA Bind-n-Seq (RBNS) analysis ^26^ with a random 20-mer RNA library and lysates of 293FT cells expressing FLAG–tagged Bicc1 (Fig. 6a). Analysis of the RBNS data set showed that the GAC motif was highly enriched by interaction of Bicc1 with the RNA library (Fig. 6b and 6c). We also found that YGAC (where Y represents a pyrimidine) and GACR (where R represents a purine) motifs possessed higher enrichment scores than did other GAC motifs (Fig. 6d), suggesting that Bicc1 binds preferentially to YGAC and GACR motifs. Metagene analysis with the 200-nucleotide proximal region of the 3’-UTR of mouse mRNAs revealed that the frequency of GACR and GAC was significantly enriched in that of *Dand5* mRNA (Fig. 6e and 6f). Scanning for GAC-containing motifs over the full-length 3’-UTR of *Dand5* mRNAs of mammals and *Xenopus* (Fig. 6g) revealed that their distribution was not uniform. In particular, the GACR motif was highly enriched in the 200-nucleotide proximal region. Importantly, YGAC/GACR motifs were also abundant in the 200-nucleotide region of the 3’-UTR of *Xenopus Dand5* mRNA (Fig. 6g), even though there was no marked sequence conservation between the 3’-UTR of mammalian *Dand5* mRNAs and that of *Xenopus Dand5* mRNA. Taken together, these results suggest that the 200-nucleotide proximal region of the 3’-UTR, which contains multiple Bicc1 binding motifs, mediates the binding of *Dand5* mRNA to the KH2 domain of Bicc1. However, Bicc1-dependent down-regulation of *Dand5* mRNA at the node of the mouse embryo in response to nodal flow likely requires additional factors that are absent or inactive in HEK293T cells.

**Figure 6.**
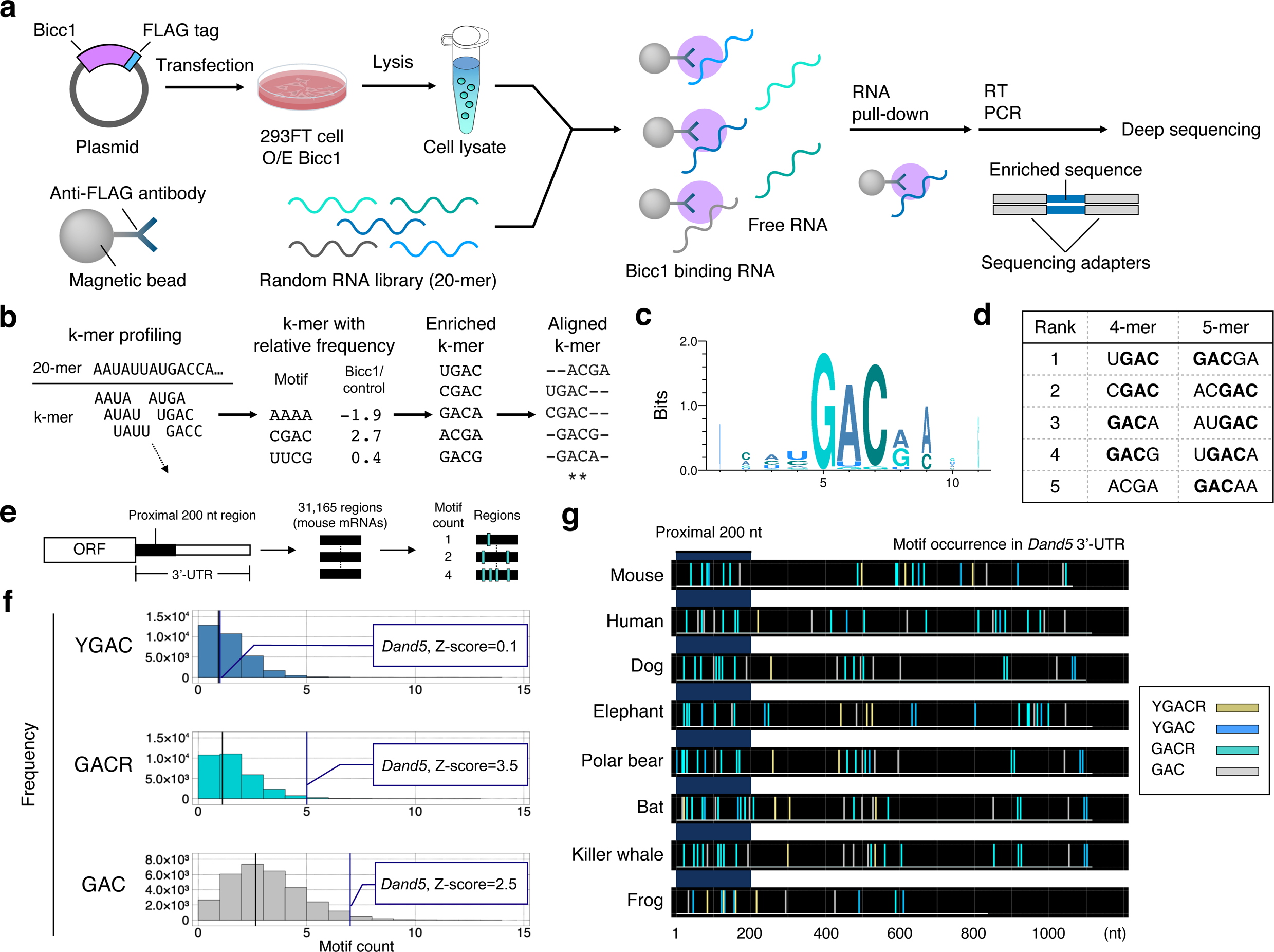
Determination of Bicc1 Binding Motifs in RNA **a** Schematic representation of RNA Bind-n-Seq (RBNS), which determines RNA motifs enriched by target proteins with the use of a random RNA sequence library. 293FT cells were transfected with a plasmid for overexpression (O/E) of FLAG-tagged Bicc1. Cell lysates containing the Bicc1-FLAG protein were then mixed with a random RNA sequence library, and resulting RNA-protein complexes were isolated by immunoprecipitation with magnetic bead–conjugated antibodies to FLAG. Finally, the isolated RNA sequences were converted to a DNA library by RT-PCR for deep sequencing. **b** Analysis of the RBNS data set. The number of each k-mer (where k = 4, 5, or 6) RNA sequence was compared between cells transfected with the Bicc1-FLAG expression plasmid and those subjected to mock transfection (control). **c** A motif logo generated from aligned hexamers significantly enriched by Bicc1-FLAG. **d** Enriched 4-mer and 5-mer sequences sorted by relative frequency determined by comparison of the Bicc1-FLAG and control RBNS data. **e** Schematic representation of metagene analysis for the 200-nucleotide proximal region of the 3’-UTR of mouse mRNAs. A total of 31,16<u>5</u> regions extracted from mouse genes (mm10) was searched with the indicated target motifs. **f** Histogram of motif frequency revealed by metagene analysis. The vertical black and blue lines indicate the averaged frequency of each target motif and the frequency of each target motif in the 200-nucleotide proximal region of the 3’-UTR of *Dand5* mRNA. **g** Maps of GAC-containing motifs in the 3’-UTR of *Dand5* mRNAs for the indicated species.

### Decay of *Dand5* mRNA on the Left Side of the Node Requires the Ccr4-Not Deadenylase Complex

In *Drosophila*, an ortholog of Bicc1 regulates mRNA decay by recruiting the Ccr4-Not complex through direct binding to its Not3/5 subunit ^14^. Although Bicc1-mediated mRNA decay has not been observed previously ^10, 27^, several Ccr4-Not subunits were also found to be enriched by mouse Bicc1 in a protein interactome screen in T-Rex HEK293 cells ^28^. To validate this interaction and to assess whether it might involve Bicc1 polymerization or RNA binding, we first performed co-immunoprecipitation assays in HEK293T cells. Both endogenous Cnot1 and FLAG-tagged Cnot3 co-immunoprecipitated with HA-Bicc1 as well as with a mutant derivative (mutD) that is unable to undergo polymerization as a result of disruption of the SAM-SAM binding interface (Figure 7A). Similarly, deletion of the entire SAM domain (ΔSAM) or of the three KH domains (ΔKH) of Bicc1 did not inhibit Cnot1 or Cnot3-FLAG binding (Fig. 7a), suggesting that the interaction is mediated by the region of Bicc1 located between the SAM and KH domains. Immunofluorescence staining revealed a nonuniform distribution of Cnot3-FLAG in the cytoplasm, with the distribution overlapping in part with that of HA-Bicc1 foci. In addition, colocalization with discrete Cnot3-FLAG foci was also observed with HA-Bicc1(mutD), confirming that Cnot3 binding to Bicc1 is independent of Bicc1 polymerization (Fig. 7b).

**Figure 7.**
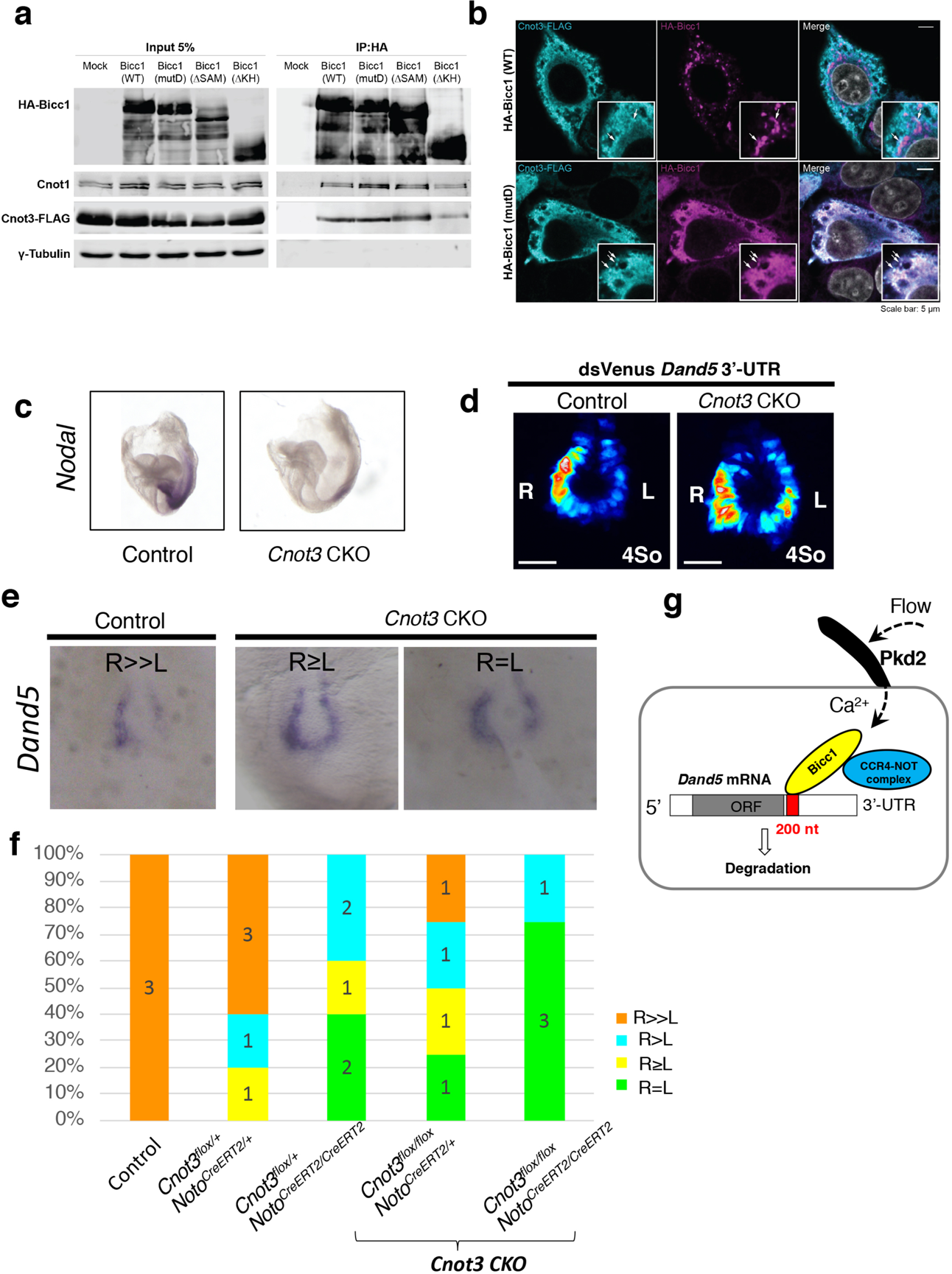
Decay of *Dand5* mRNA on the Left Side of the Node Requires the Ccr4-Not Deadenylase Complex **a** Extracts of HEK293T cells transfected with expression plasmids for Cnot3-FLAG and either WT or the indicated mutant forms of HA-Bicc1 (or with the corresponding empty vector, Mock) were subjected to immunoprecipitation with antibodies to HA. The resulting immunoprecipitates as well as a portion (5%) of the original cell extracts were then subjected to immunoblot analysis with antibodies against HA, Cnot1, FLAG, or γ-tubulin (loading control). **b** Immunofluorescence staining of Cnot3-FLAG and HA-Bicc1 (WT or mutD) in transfected HeLa cells. Arrows in magnified areas (insets) indicate overlap of fluorescence signals. Scale bars, 5 µm. **c** WISH analysis of *Nodal* expression in control and *Cnot3* CKO embryos at E8.0. Note that *Nodal* expression in LPM was lost in the mutant embryo. **d** Fluorescence of dsVenus in control and *Cnot3* CKO embryos harboring the *NDE-Hsp-dsVenus-3’-UTR* transgene at the four-somite stage. Note that Venus fluorescence is increased on the left side of the *Cnot3* CKO embryo. Scale bars, 50 µm. **e** WISH analysis of *Dand5* mRNA in control and *Cnot3* CKO embryos at five- to six-somite stages. The R > L asymmetry of *Dand5* mRNA in the control embryo is attenuated or lost in the *Cnot3* CKO embryos. **f** Summary of the L-R pattern of *Dand5* mRNA at the node of embryos of the indicated genotypes as in (E). **g** Model for the degradation of *Dand5* mRNA in crown cells of the node in response to nodal flow. See text for details.

Finally, we examined whether Cnot3 contributes to the decay of *Dand5* mRNA in crown cells at the node. A Cnot3 conditional knockout (CKO) mouse was generated by crossing a *Cnot3*^flox/flox^ mouse ^29^ with a *Noto*^CreERT2/+^ mouse ^30^, the latter of which expresses Cre recombinase in a tamoxifen-inducible manner specifically at the node of the embryo beginning at E8.0. Control (*Cnot3*^flox/+^;*Noto*^CreERT2/+^) and CKO (*Cnot3*^flox/flox^;*Noto*^CreERT2/+^) embryos isolated from tamoxifen-treated dams were first examined for *Nodal* expression, which was found to be maintained at the node but was lost in LPM of the *Cnot3* CKO embryos (Fig. 7c), consistent with the possibility that the stability of *Dand5* mRNA in crown cells was increased. We then examined *Dand5* expression in control and CKO embryos at the five- to six-somite stages. L-R asymmetry of *Dand5* mRNA was lost or attenuated in the CKO embryos (Fig. 7d, e and f), suggesting that the Ccr4-Not complex in crown cells indeed plays a role in the degradation of *Dand5* mRNA in response to nodal flow.

## DISCUSSION

To delineate how fluid flow establishes left-right asymmetry in the mouse embryo, we have generated a Venus transgene that reports the decay of *Dand5* mRNA in crown cells of the node and its acceleration by directional fluid flow and by Pkd2 specifically on the left side. Our data show that a proximal 3’-UTR fragment mediating asymmetric *Dand5* mRNA expression binds and genetically interacts with the RNA binding protein

Bicc1 and is enriched for the newly identified Bicc1 consensus recognition motif GAC. In turn, Bicc1 binds to the Ccr4-Not deadenylase complex which mediates increased degradation of the *Dand5* mRNA in response to Pkd2 stimulation by leftward fluid flow.

Bicc1 binds to various target mRNAs, but no consensus motif recognized by this protein has been identified. Our comparison of various mammalian *Dand5* genes revealed that conservation among the 3’-UTR sequences is largely limited to the 200-nucleotide proximal-most region which we found to be required for establishment of L-R asymmetry of *Dand5* mRNA in crown cells. Strikingly, no sequence element appears to be highly conserved in this region or elsewhere in the 3’-UTR among vertebrate *Dand5* genes. However, our RBNS analysis revealed a binding preference of Bicc1 for GACR and YGAC motifs, which are enriched in the 200-nucleotide proximal region of the 3’-UTR of mammalian *Dand5* mRNAs. Given that none of these motifs or their flanking sequences are absolutely conserved, they may act in a combinatorial manner to mediate the binding of Bicc1 oligomers. However, testing of their relative contributions by mutagenesis would be complicated by likely effects on mRNA folding. Individual KH domains bind to 4-nucleotide motifs, where nucleobase recognition is mediated primarily by positions 2 to 4^22^. We found that association of Bicc1 with the Dand5 3’-UTR required the KH2, whereas KH1 may increase the binding affinity. Therefore, and since GACR and YGAC emerged from our RBNS screen as the predominant consensus motifs, it is possible that all three Bicc1 KH domains recognize the same consensus motif but with differential affinities, or that only KH2 has a sufficiently high affinity for RNA to bind a single motif on its own. In keeping with either of these scenarios, a 32-nucleotide region of *Xenopus Cripto1* mRNA that has previously been shown to bind Bicc1 contains an AGACA motif that fits both NGAC (where N represents any nucleotide) and GACR consensus sequences ^31^. Furthermore, a 36-nucleotide region of the 3’-UTR of *Xenopus Dand5S* mRNA required for its flow induced decay at the left-right organizer contains three YGAC sequences, with two YGAC motifs also being present in a nearby region (Maerker et al., accompanying paper). Given that mRNAs form stem-loop structures, Bicc1 may recognize YGAC and GACR motifs that are present in the accessible regions of the secondary structure, with the presence of multiple GAC motifs possibly allowing synergistic binding to several different regions.

Bicc1 targets mRNAs such as those of mouse *Adcy6* or *Xenopus Cripto1* to inhibit their translation ^10, 27, 32^. In crown cells of the mouse node as well as in ciliated cells at the gastrocoel roof plate of *Xenopus* embryos, however, the level of *Dand5* mRNA is asymmetric (R > L). Furthermore, *Dand5* mRNA in crown cells on the left side of the mouse node decreases in abundance in response to nodal flow ^8^, suggestive of increased mRNA decay. Bicc1-mediated regulation of certain mRNAs has been shown to involve microRNAs (miRNAs) ^10, 33^, and miRNA-mediated inhibition of target mRNAs typically involves both translational repression and mRNA decay ^34^ mediated by Ccr4-Not ^35^. Consistent with a potential role for miRNAs in the regulation of *Dand5* mRNA in crown cells of the mouse embryo, deletion of *Dicer* in these cells abolished asymmetric *Nodal* expression in the lateral plate (Maerker et al., accompanying paper). However, given that no miRNA seed sequences are highly conserved in the 3’-UTR of mammalian *Dand5* mRNAs (Figure 3A) and that Ccr4-Not can also be recruited by specific RNA-binding proteins independently of miRNAs ^36^, the regulation of *Dand5* mRNA by Bicc1 may be mediated by an miRNA-independent pathway. In all, our data indicate that inhibition of Dand5 expression at the mouse node is dependent on the 3’-UTR of the mRNA and mediated by Bicc1 and Cnot3 at the level of mRNA decay.

The degradation of *Dand5* mRNA in crown cells occurs predominantly on the left side of the mouse node. How can the Bicc1-Cnot3 pathway be activated on the left side but not on the right side? Several lines of evidence implicate Ca^2+^ in the decay of *Dand5* mRNA. First, the Dand5-dependent asymmetry (R < L) of Nodal activity in crown cells is disrupted by various Ca^2+^ blockers including thapsigargin ^5^ and ionophore^37^. Furthermore, R > L asymmetric expression of the *NDE-Hsp-dsVenus-3’-UTR* transgene at the node in the present study was impaired by thapsigargin treatment and by the absence of Pkd2. The decay of *Dand5* mRNA may therefore be induced by the Ca^2+^ influx triggered by sensing of nodal flow by the immotile cilia of crown cells. The resulting increase in the cytosolic concentration of Ca^2+^ may influence, directly or indirectly, either the interaction between components of the pathway responsible for mRNA decay (such as between Bicc1 and *Dand5* mRNA, or between Bicc1 and Ccr4-Not) or the deadenylase activity of Ccr4-Not. Of note in this regard, crown cells at the mouse node express calmodulin and proteins such as Inversin with a calmodulin binding motif, some of which are known to be essential for L-R asymmetry ^38, 39^. Future studies are thus warranted to probe further the role of Ca^2+^ in the regulation of *Dand5* mRNA decay.

## METHODS

### Mouse Strains

All mouse experiments were performed in accordance with guidelines of the RIKEN Center for Biosystems Dynamics Research and under an institutional license (A2016-01-6). Mice were maintained in the animal facility of the RIKEN Center for Biosystems Dynamics Research. *Noto-Cre*^ERT2^ mice ^30^, *Cnot3*^flox/+^ mice ^29^, *iv* mice ^18^, and *Pkd2* mutant mice ^3^ were described previously. Expression of the *Noto-Cre*^ERT2^ transgene in embryos was induced by oral administration of tamoxifen (Sigma) in corn oil to pregnant mice at a dose of 5 mg both 24 and 12 h before the late headfold stage.

### Cell Lines

HEK293T and HeLa cells were cultured in DMEM (Sigma) supplemented with 10% FBS (Sigma), 1% GlutaMAX (Thermo Fisher Scientific), and 1% gentamicin (Thermo Fisher Scientific). The cells were transfected with expression plasmids with the use of jetPEI (Polyplus Transfection). IMCD cells were cultured in DMED/F12 (1:1) (Gibco 11320-033) with 10 % FBS. For RNBS analysis, the human embryonic kidney cell line 293FT (Invitrogen, #R70007) was cultured under a humidified atmosphere of 5% CO_2_ at 37°C in Dulbecco’s modified Eagle’s medium (DMEM) (Nacalai tesque, #08459-64) supplemented with 10% fetal bovine serum (FBS) (Biosera, #554-02155, lot #10259), 2 mM L-glutamine (Thermo Fisher Scientific, #25030-081), 1× MEM Nonessential Amino Acids (Thermo Fisher Scientific, #11140-050), and 1 mM sodium pyruvate (Sigma, #S8636-100ML).

### Transgene Design

For generation of a transgene (*NDE-Hsp-dsVenus-3’-UTR*) that confers expression of dsVenus at the node of mouse embryos, the ORF for dsVenus was linked to the 1.2-kb DNA sequence for the 3’-UTR of mouse *Dand5* mRNA and placed under the control of four copies of the crown cell–specific enhancer (NDE) of the mouse *Nodal* gene and the mouse *Hsp68* promoter. A corresponding transgene lacking the 200-bp proximal region of the 3’-UTR sequence (*NDE-Hsp-dsVenus-3’-UTRΔ200*) was also generated.

### Generation of Transgenic and Mutant Mice

*Dand5*^Δ200/Δ200^ embryos (Figure S1) and *Bicc1*^Δex1/ Δex1^ embryos (Figure S2) were generated with the use of the CRISPR/Cas9 system. For the generation of transgenic mice, each transgene was microinjected into the pronucleus of fertilized eggs obtained by crossing C57BL/6J female and male mice. The eggs were then transferred to pseudopregnant mothers. In some instances (embryos shown in Figure 3B, D, E), transgenic embryos were directly examined before establishment of stable transgenic lines. *Dand5* and *Bicc1* mutant mice and embryos were genotyped by genomic PCR analysis with the primers indicated in Supplementary Figures 1 and 2.

### Plasmids, Cloning, and Mutagenesis

The plasmid pCMV6-Entry::Cnot3-Myc-Flag encoding mouse Cnot3 was obtained from Origene (MR210463). The pCMV-SPORT6::HA-Bicc1 plasmids for WT, ΔKH, ΔSAM, and mutD forms of mouse Bicc1 were previously described (Maisonneuve et al., 2009; Rothé et al., 2015). Vectors encoding mutKH1, mutKH2, mutKH3, or mutKH1,2,3 were generated by overlap-extension PCR and subcloning between the SalI and BstBI sites of pCMV-SPORT6::HA-Bicc1. The sequences of all mutated constructs were verified by Sanger sequencing.

### WISH Analysis

WISH was performed according to standard procedures with digoxigenin-labeled riboprobes specific for *Dand5*, *Nodal*, and *dsVenus* mRNAs ^40^.

### Immunofluorescence Analysis

Dissected embryos were fixed with 4% paraformaldehyde, dehydrated with methanol, and permeabilized with phosphate-buffered saline (PBS) containing 0.1% Triton X-100. They were incubated overnight at 4°C with primary antibodies at the following dilutions: anti-Bicc1 (1:100 dilution, rabbit polyclonal, Sigma), anti–acetylated tubulin (1:100 dilution, mouse monoclonal, Sigma), anti-ZO1 (1:10 dilution, mouse monoclonal, clone ZO1-1A12, Invitrogen), and anti-Odf2 (1:100 dilution, rabbit polyclonal, Abcam). The embryos were washed with PBS containing 0.1% Triton X-100 before exposure to Alexa Fluor–conjugated secondary antibodies (Invitrogen) and DAPI (Wako). The node of stained embryos was excised, placed on a slide glass with silicone rubber spacers, covered with a cover glass, and imaged with an Olympus FV1000 or FV3000 confocal microscope. For immunostaining of HeLa cells, cells cultured in six-well plates and transfected with 1 µg of each expression vector for 24 h were transferred to sterile coverslips in the wells of a 24-well plate. After culture for an additional 24 h, the cells were fixed for 10 min at –20°C in methanol, washed with PBS, and incubated at room temperature first for 1 h in PBS containing 1% bovine serum albumin as a blocking agent and then for 2 h in the blocking solution containing antibodies to HA (1:500, rabbit monoclonal, Sigma) and to FLAG (1:500, mouse monoclonal clone M2, Sigma). After washing with PBS, the cells were incubated for 1 h at room temperature with Alexa Fluor 568– or Alexa Fluor 647–conjugated secondary antibodies to mouse or rabbit immunoglobulin G, respectively, in blocking buffer containing DAPI (1:10,000 dilution). Images were acquired with a Zeiss LSM700 confocal microscope.

### Observation of Nodal Flow by PIV Analysis

Nodal flow was observed by multipoint scanning confocal microscopy. Particle image velocimetry (PIV) analysis was performed as described previously ^17^. Recovered embryos were first cultured under 5% CO_2_ for 30 min at 37°C in DMEM supplemented with 75% rat serum. The region containing the node was then excised, and the node cavity was filled with DMEM supplemented with 10% FBS and 0.2µm-diameter fluorescent microbeads (Invitrogen, F8811). The motion of the beads was monitored for 10 s in planes of +5 and +10 µm relative to the cavity (21 frames per second) with the use of a CSU-W1 confocal unit (Yokogawa) and an iXon-Ultra EMCCD camera (Andor Technology) connected to an IX83 microscope (Olympus) fitted with a ×60 objective lens. Time-series images for PIV analysis were captured at a resolution of 512 by 512 pixels and were processed with interrogation windows of 16 by 16 pixels with 50% overlap, corresponding to a spatial resolution of 4.3 by 4.3 mm. The time-averaged velocity distributions were calculated for 10-s intervals.

### Quantitative Analysis of Basal Body Position

The average basal body position (ABP) representing the relative position of the basal body in each node cell along the A-P axis (vertical) was analyzed as described previously ^41, 42^. Confocal images of the node stained with antibodies to Odf2 and to ZO-1 were obtained to determine the position of the basal body in each node cell. For characterization of the shape and orientation of the node cells, the outline of each cell was calculated from the pattern of ZO-1 staining with the use of an ImageJ software plugin (http://rsb.info.nih.gov/nih-image) to apply watershed segmentation. The basal body was traced manually according to the Odf2 staining image, and the *x* and *y* coordinates of each basal body were recorded with the use of the graphical user interface (GUI) of MATLAB software. The relative value for the position of the basal body in each cell was calculated from the coordinate data with the anterior and posterior ends of the cell expressed as –1.0 and +1.0, respectively.

### Mouse Embryo Culture

Embryos were collected at E7.5. Those at the early headfold stage were selected and cultured with rotation under 5% CO_2_ at 37°C in DMEM supplemented with 75% rat serum. Where indicated, thapsigargin was added to the medium at a final concentration of 500 nM for 1 h and the embryos were then cultured in the absence of the drug.

Embryos were cultured under conditions of artificial fluid flow as described previously^19^. The node cavity was imaged with an Olympus FV1000 confocal microscope.

### dsVenus Fluorescnce Imageing

The embryonic region containing the node of the mouse embryo was excised in DMEM supplemented with 75% rat serum, placed on slide glasses with silicone rubber spacers, covered with cover glasses, and imaged with Olympus FV1000 confocal microscope.

### Protein Co-immunoprecipitation and Western blot Analysis

HEK293T cells in two 10-cm dishes per condition (10^7^ cells per dish) were transfected with expression plasmids for Cnot3-FLAG (2 µg) and either HA-Bicc1(WT) (2 µg), HA-Bicc1(ΔKH) (1 µg), HA-Bicc1(ΔSAM) (4 µg), or HA-Bicc1(mutD) (4 µg). After incubation for 3 days, cells from the two dishes for each condition were then washed with PBS and pooled in extraction buffer containing 20 mM Tris-HCl (pH 7.4), 2.5 mM MgCl_2_, 100 mM NaCl, 5% glycerol, 1 mM dithiothreitol (DTT), 0.05% Nonidet P-40 (NP-40), RNasin (Promega), phosphatase inhibitors (Sigma), and protease inhibitors (Roche). Total cell extracts were prepared by ultrasonication followed by two rounds of centrifugation at 10,000 × *g* for 5 min at 4°C to remove debris. Supernatants were incubated with 20 µl of anti-HA beads (Sigma) for 2.5 h at 4°C on a rotating wheel. The beads were rinsed four times for 10 min in wash buffer (20 mM Tris-HCl pH 7.4, 2 mM MgCl_2_, 200 mM NaCl, 1 mM DTT, and 0.1% NP-40), suspended in Laemmli buffer, and directly loaded on reducing SDS PAGE gels to size-separate eluted proteins. For immunoblot analysis, proteins were transferred to nitrocellulose membranes, blocked with skim milk and incubated over night with the antibodies indicated, and with fluorescently labelled secondary antibodies for analysis on a Odyssey CLx scanner (LI-COR Biosciences).

### RNA Co-immunoprecipitation in HEK293T Cells

HEK293T cells were transfected as described above, but with an expression plasmid for the ORF of dsVenus linked to the WT or *Δ200* forms of the 3’-UTR sequence of mouse *Dand5* mRNA (2 µg per dish) together with a plasmid encoding WT or ΔKH mutant forms of HA-Bicc1 (2 or 1 µg per dish, respectively). Cytoplasmic extracts prepared after 36 h as described above were passaged eight times through a syringe needle (no. 30) followed by centrifugation twice at 10,000 × *g* for 5 min at 4°C. After setting aside 5% of each supernatant as input, the remainder was immunoprecipitated as described above. While a portion of the beads (10%) was analyzed by immunoblotting as described above, the remainder of the beads and half of the input samples were subjected to phenol-chloroform extraction. After ethanol precipitation isolated RNA was treated with RQ1 DNase (Promega) and converted to cDNA by PrimeScript Reverse Transcriptase Kit (Takara). The resulting cDNA was then subjected to PCR or qPCR using Phire Green Hot Start II PCR Master Mix (Promega) or GoTaq qPCR Master Mix (Promega), respectively, with the following primers: F1 (GCTGAGCATCCTAGAGGAATGC), F2 (TGCCACAATCACTAACTCACGTC, and R (TGTCTTGGACACTGGGACGC) for the 3’-UTR of *Dand5* mRNA, and ACAGAGCCTCGCCTTTGCC and CTCCATGCCCAGGAAGGAAGG (forward and reverse, respectively) for β-actin mRNA. The amount of co-immunoprecipitated mRNA as a percentage of the input was calculated with the formula: 100 × 2[(Ct(Input) − log_2_(100/2.5) − Ct(IP)], where the value 2.5 represents 2.5% of the original cytoplasmic extract, and where Ct is the cycle threshold for qPCRs on input or immunoprecipitate (IP) samples. Fold enrichment was calculated relative to cells transfected with the corresponding empty vector for HA-Bicc1.

### RNA Co-immunoprecipitation in IMCD3 Cells

Mouse IMCD3 cells stably transfected with an expression plasmid for the ORF of dsVenus linked to the WT or *Δ200* forms of the 3’-UTR sequence of mouse *Dand5* mRNA (3 × 10^6^ cells in each of two 10-cm dishes per condition) were washed with PBS, collected by centrifugation, and suspended in 400µl of a hypotonic buffer containing 10 mM Tris-HCl (pH 7.4), 1.5 mM MgCl_2_, 10 mM NaCl, 1 mM DTT, and 0.1% NP-40. The cell suspension was centrifuged at 400 × *g* for 5 min to remove nuclei and debris, and the resulting supernatant (cytoplasmic fraction) was mixed with an equal volume of the extraction buffer (30 mM Tris-HCl, pH 7.4, 3.5 mM MgCl_2_, 190 mM NaCl, 10% glycerol, 1 mM DTT, Phosphatase inhibitor and protease inhibitors). Protein G–Sepharose beads that had been incubated with antibodies to Bicc1 (1:25 dilution, Sigma) for 2 h at 4°C were then added to each cytoplasmic extract and incubated for 2.5 h at 4℃. The beads were collected by centrifugation and washed three time for 10 min with the wash buffer described in the previous section, after which RNA was isolated from the washed beads by phenol-chloroform extraction followed by ethanol precipitation and DNase treatment as described above. The isolated RNA was subjected to RT with SuperScript IV VILO reverse transcriptase (Thermo Fisher Scientific), and the resulting cDNA was subjected to qPCR with primers (forward and reverse, respectively) specific for dsVenus (5’-CGACCACTACCAGCAGAACA-3 and 5’-GAACTCCAGCAGGACCATGT-3’) or β-actin (5’-GCTACAGCTTCACCACCACA-3’ and 5’-TCTCCAGGGAGGAAGAGGAT-3’).

### RBNS Analysis

RBNS analysis was performed essentially as described by others ^26^. A double-stranded DNA template for in vitro transcription of a 20-mer random RNA library was synthesized with a primer extension reaction in which 100 µl of a reaction mixture containing 1× Platinum SuperFi PCR Master Mix (Thermo Fisher Scientific, #12358-010), 1× SuperFi GC Enhancer (Thermo Fisher Scientific), 100 nM DNA template oligomer (5’-GAAATTAATACGACTCACTATAGGACGTGACACGACGTGCGCN_20_GCGTACG TCGGACCTCAGGTCGACCATGGACGC-3’, where N_20_ is the DNA sequence encoding the 20-mer RNA sequence), 100 nM primer (5’-GCGTCCATGGTCGACCTGAGGTCC-3’), and nuclease-free water was incubated at 98°C for 130 s, at 50°C for 2 min, and then at 72°C for 10 min. The synthesized DNA template was purified with the use of a Monarch PCR & DNA Cleanup Kit (New England Biolabs, #T1030L). The random RNA library was then transcribed with the use of a MEGAshortscript T7 Transcription Kit (Thermo Fisher Scientific, #AM1354) in a reaction mixture containing 1× Reaction Buffer, 1× T7 Enzyme Mix, 7.5 mM ATP, 7.5 mM UTP, 7.5 mM GTP, 7.5 mM CTP, and 9.15 pmol of the DNA template. The mixture was incubated at 37°C for 6 h, after which the DNA template was digested with TURBO DNase for 30 min at 37℃ and the transcribed RNA library (5’-GGACGUGACACGACGUGCGCN_20_GCGUACGUCGGACCUCAGGUCGACCAUGGACGC-3’) was purified with an RNA Clean & Concentrator (Zymo Research, #R1016).

For generation of an expression vector for Bicc1-FLAG (pcDNA3.1-Bicc1-FLAG), the Bicc1 ORF was first amplified from Bicc1/mBS and then inserted between the BamHI and NotI sites of pcDNA3.1/myc-His A (Invitrogen, #V80020) to generate pcDNA3.1-Bicc1-myc-His A. The sequences corresponding to the c-Myc and His tags in pcDNA3.1-Bicc1-myc-His A were replaced with that for the FLAG tag by PCR-based mutagenesis, thereby generating pcDNA3.1-Bicc1-FLAG. 293FT cells (3 × 10^6^ cells) were seeded in a 10-cm cell culture dish and cultured for 24 h before transfection for 24 h with 15 µg of pcDNA3.1-Bicc1-FLAG with the use of Lipofectamine 3000 (Thermo Fisher Scientific, #L3000008) according to the manufacturer’s instructions. As a mock transfection control, cells were treated with Lipofectamine 3000 alone. The cells were then collected by centrifugation, washed with ice-cold PBS, and lysed by incubation for 30 min on ice, with intermittent vortex mixing, in 1 ml of Cell Lysis Buffer (Invitrogen, #FNN0021) supplemented with 1× cOmplete Protease Inhibitor Cocktail (Roche, #04693116001) and 1 mM phenylmethylsulfonyl fluoride (Thermo Fisher Scientific, #36978). The cell lysates were centrifuged at <u>13,000 rpm</u> for 10 min at 4°C, and the resulting supernatants were collected, <u>assayed for protein concentration with the use of a Pierce BCA Protein Assay Kit (Thermo Fisher Scientific, #23227),</u> adjusted to a protein concentration of 700 µg/ml with cell lysis buffer, and stored at – 80°C.

The cell lysate containing Bicc1-FLAG was diluted fourfold with the mock cell lysate for immunoprecipitation. For preparation of antibody-coated magnetic beads, 50µl of Dynabeads Protein G (Thermo Fisher Scientific, #DB10003) were conjugated with 10 µg of M2 mouse monoclonal antibodies to FLAG (Sigma, #F1804) according to the manufacturer’s instructions. The antibody-coated beads were suspended in 400 µl of Ab Binding & Washing Buffer [PBS containing 0.02% Tween-20 (Roche, #11332465001)] and mixed with 400 µl of the cell lysate. After incubation for 1 h at 4°C with rotation, the beads were separated with a magnet and the supernatant removed. The beads were washed with 1 ml of RNP Binding Buffer [20 mM Tris-HCl (pH 7.4), 2.5 mM MgCl_2_, 80 mM NaCl, 20 mM KCl, 5% glycerol (Sigma, #G5516-100ML), 1 mM DTT (Thermo Fisher Scientific, #A39255), 0.05% NP-40 (Thermo Fisher Scientific, #85124)], suspended in 500 µl of RNP Binding Buffer containing 500 pmol of the random RNA library, incubated for 16 h at 4°C with rotation, isolated with the magnet, and washed five times with 1 ml of RNP Binding Buffer. The beads were then incubated for 3 min at 95°C in 200 µl of an elution buffer (1% SDS, 10 mM Tris-HCl, and 2 mM EDTA) before separation with the magnet, and the supernatant was collected. <u>T</u>he eluted RNA was isolated by phenol-chloroform extraction and ethanol precipitation. The RNA pellet was suspended in 24 µl of nuclease-free water and subjected to RT with SuperScript IV Reverse Transcriptase (Thermo Fisher Scientific, #18090010). Prior to reverse transcription, 20 µL of purified RNA library was mixed with 1 µL of 10 µM reverse primer (5’-GCGTCCATGGTCGACCTGAGGTCC-3’), 1 µL of 10 mM dNTPs, and 4µL of nuclease water, and then denatured at 65°C for 5 minutes. The mixture was combined with a premix of 8 µL of 5X SSIV Buffer, 2 µL of 100 mM DTT, 2 µL of RNase OUT (Thermo Fisher Scientific, #10777019), and 2 µL of SuperScript IV

Reverse Transcriptase. The mixture was incubated at 50°C for 10 minutes, then at 85°C for 10 minutes. To prepare an input random RNA library, 0.5 pmol of input random RNA library was also reverse transcribed. After digestion of RNA with 2 µl of RNaseH for 20 min at 37°C, the cDNA preparation was amplified by PCR in a reaction mixture containing 1× Platinum SuperFi PCR Master Mix, 5 µM forward primer (5’-TCGTCGGCAGCGTCAGATGTGTATAAGAGACAGGGACGTGACACGACGTGC-3’), 5 µM reverse primer (5’-GTCTCGTGGGCTCGGAGATGTGTATAAGAGACAGGGCGTCCATGGTCGACCTGAGGTCCGACG-3’), 1× SuperFi GC Enhancer, and 2 µl of cDNA and adjusted to 25 µl with nuclease-free water. The incubation protocol included an initial denaturation at 98°C for 30 s, 20 cycles of denaturation at 98°C for 10 s and extension at 72°C for 10s, and a final extension at 72°C for 5 min. Single-stranded DNA was digested with Illustra ExoProStar exonuclease I (GE Healthcare Life Sciences, #US78211), and the PCR products were purified with the use of a MinElute PCR Purification Kit (Qiagen, #28006) before the addition of Illumina sequencing barcodes by 10 cycles of PCR with index primers from a Nextera XT Index Kit (Illumina, #FC-131-1001). The reaction mixture contained 1× Platinum SuperFi PCR Master Mix, 5 µl of Index 1 (i7) primer, 5µl of Index 2 (i5) primer, 1× SuperFi GC Enhancer, and at most 5 µl of the PCR products, and the volume was adjusted to 50 µl with nuclease-free water. The incubation protocol included an initial denaturation at 98°C for 30 s; 10 cycles of denaturation at 98°C for 10 s, annealing at 55°C for 10 s, and extension at 72°C for 20 s; and a final extension at 72°C for 5 min. The resulting sequencing library was quantified with the use of a KAPA Library Quantification Kit (Illumina) and Universal qPCR Mix (Kapa Biosystems, #07960140001). The library was sequenced for 150 cycles on the Illumina MiSeq platform and with a MiSeq Reagent Kit v3 (Illumina, #MS-102-3001). FASTQ files were processed as follows. The adapter sequences of the reads were trimmed by cutadapt 1.10 with parameter -e set to 0.2. Low-quality reads (averaged quality score of <30) were filtered with a custom script written in Julia 1.1. The filtered reads were split into overlapping k-mers (k = 4, 5, or 6); for example, the 20-mer sequence yielded 15 hexamers. The frequency of each k-mer was determined and normalized by the sum of all frequencies. With *k* denoting each k-mer, the following formula defines relative frequency: relative frequency(*k*) = normalized frequency(*k*)_Bicc1_/normalized frequency(*k*) _Control_. To identify Bicc1-bound sequence elements, we collected the hexamers whose Z-score for the relative frequency was >3.0. The enriched hexamers were aligned by Clustal Omega with default parameters. The motif logo was generated from the aligned hexamers by Weblogo3.

### Metagene analysis

Mouse 3’-UTR sequences (mm10) were obtained from the UCSC Table Browser by specifying the track as ALL GENCODE V22 and the table as Basic. The 200-nucleotide proximal regions were extracted, and duplicated sequences were removed. The number of motifs present in each region was then counted.

### QUANTIFICATION AND STATISTICAL ANALYSIS

In Figure 5 and Supplementary Figure 2, the statistical analyses used the two-tailed Student’s t-test with *p < 0.05, **p < 0.01, ***p < 0.005 taken as significance. In Figure 6, StatsBase package with Julia 1.1 was used to calculate Z-scores.

### DATA AND CODE AVAILABILITY

The GEO accession number for the RBNS sequencing data reported in this paper is GSE140931 (token for review: wpelesgyxvydzqh). The codes used in this study are available from the corresponding authors upon request. All other relevant data supporting the key findings of this paper are available from the corresponding authors upon request.

## ACKNOWLEDGMENTS

We thank K. Okamoto (University of Tokyo) for PIV analysis software; Y. Igarashi for the software for quantitative analysis of basal body position; H. Sasaki for *Noto*^CreERT2/+^ mice; and M. Yamagata and S. Wada for technical support and designing oligonucleotides for RBNS analysis, respectively. This study was supported by grants from the Ministry of Education, Culture, Sports, Science, and Technology (MEXT) of Japan (no. 17H01435) and from Core Research for Evolutional Science and Technology (CREST) of the Japan Science and Technology Agency (no. JPMJCR13W5) to H.H.; by a Grant-in-Aid (no. 18K14725) for Early-Career Scientists from the Japan Society for the Promotion of Science (JSPS), a Kakehashi grant from BDR-Otsuka Pharmaceutical Collaboration Center, and a research grant (no. 2018M-018) from the Kato Memorial Bioscience Foundation to K.M.; by grants from the NIBB Individual Collaborative Research Program (nos. 16-312 and 17-316) and a Sinergia grant (no. CRSI33_130662) from the Swiss National Science Foundation to D.B.C.; and by a KAKENHI grant (no. 15H05722) from JSPS and a research grant from The Mitsubishi Foundation to H.S.

## AUTHOR CONTRIBUTIONS

K.M. performed most experiments with mouse embryos. Y.I., H.N., and K.T. generated transgenic and mutant mice. B.R. performed biochemical analysis of Bicc1 and Cnot3. K.R.K., E.M., and H.O. performed RBNS analysis. K.B. and H.K. performed transgenic assays with deletion mutants. T.Y. examined the phenotype of *Cnot3* CKO mutant mice. H.H., D.B.C., H.S., K.M., B.R., and K.R.K. conceived the project and wrote the paper. All authors contributed to revision of the manuscript.

## SUPPLEMENTAL INFORMATION

**Supplementary Figure 1. Generation of Mice with the *Dand5*^Δ200^ Allele with the CRISPR/Cas9 System**

**a** Schematic representation of deletion of the 200-bp DNA sequence corresponding to the proximal-most region of the 3’-UTR of *Dand5* mRNA from the mouse genome. The red-shaded region of the WT allele was deleted to give rise to the *Dand5*^Δ200^ allele.

The positions of guide RNAs (gRNAs, green bars) and of PCR primers for genotyping (blue arrows) are indicated. **b** Nucleotide sequence of exon 2 of mouse *Dand5* showing portions related to deletion of the 200-bp fragment in (A) with the CRISPR/Cas9 system. **c** PCR-based genotyping of WT, *Dand5*^Δ200/+^, and *Dand5*^Δ200/Δ200^ mouse embryos with the indicated primers.

**Supplementary Figure 2. Generation of *Bicc1* Mutant Mice with the CRISPR/Cas9 System**

**a** *Bicc1* mutant mice were generated by CRISPR/Cas9 editing with two gRNAs (green bars) to delete exon 1. The positions of PCR primers (blue arrows) for genotyping are also indicated. **b** Relative position of the basal body in node cells of WT and *Bicc1*^Δex1/Δ ex1^ embryos at three- to five-somite stages. Each green dot corresponds to the relative position of one basal body along the anterior (A)–posterior (P) axis. The average basal body position (ABP) for each embryo is indicated by the red bar, with the corresponding value being given below each image. **c** Summary of ABP for the indicated numbers of embryos (n) determined as in (B). The p value was determined with the T test.

**Supplementary Figure 3. Binding of Endogenous Bicc1 to the *dsVenus-3’-UTR* Reporter mRNA**

**a** A cytoplasmic fraction prepared from IMCD3 cells stably expressing the *dsVenus-3’-UTR* or *dsVenus-3’-UTRΔ200* constructs was subjected to immunoprecipitation with antibodies to Bicc1, and the resulting immunoprecipitates as well as a portion of the input material were subjected to RT-qPCR analysis with PCR primers specific for dsVenus or β-actin (negative control). Data represent the ratio of co-immunoprecipitated mRNA to the input. **b** Binding of Bicc1 to the *dsVenus-3’-UTR* Reporter mRNA Does Not Affect the Abundance of the Encoded Protein. Cytoplasmic extracts of HEK293T cells transfected as in Figure 5B were subjected to immunoblot analysis with antibodies to GFP (for detection of dsVenus), to HA (for detection of HA-Bicc1), and to γ-tubulin (loading control). The antibodies to GFP detected both the intact dsVenus protein (34 kDa) and a degradation product (26 kDa).

## REFERENCES

1. Blum, M., Feistel, K., Thumberger, T. & Schweickert, A. The evolution and conservation of left-right patterning mechanisms. Development 141, 1603–1613, doi:Doi 10.1242/Dev.100560 (2014).

2. Nakamura, T. & Hamada, H. Left-right patterning: conserved and divergent mechanisms. Development 139, 3257–3262, doi:139/18/3257 [pii] 10.1242/dev.061606 (2012).

3. Pennekamp, P. et al. The ion channel polycystin-2 is required for left-right axis determination in mice. Curr Biol 12, 938–943, doi:S0960982202008692 [pii] (2002).

4. Field, S. et al. Pkd1l1 establishes left-right asymmetry and physically interacts with Pkd2. Development 138, 1131–1142, doi:dev.058149 [pii] 10.1242/dev.058149 (2011).

5. Yoshiba, S. et al. Cilia at the node of mouse embryos sense fluid flow for left-right determination via Pkd2. Science 338, 226–231, doi:10.1126/science.1222538 (2012).

6. Shiratori, H. & Hamada, H. TGFbeta signaling in establishing left-right asymmetry. Semin Cell Dev Biol 32, 80–84, doi:10.1016/j.semcdb.2014.03.029 (2014).

7. Marques, S. et al. The activity of the Nodal antagonist Cerl-2 in the mouse node is required for correct L/R body axis. Genes Dev 18, 2342–2347, doi:18/19/2342 [pii] 10.1101/gad.306504 (2004).

8. Kawasumi, A. et al. Left-right asymmetry in the level of active Nodal protein produced in the node is translated into left-right asymmetry in the lateral plate of mouse embryos. Dev Biol 353, 321–330, doi:S0012-1606(11)00158-8 [pii] 10.1016/j.ydbio.2011.03.009 (2011).

9. Maisonneuve, C. et al. Bicaudal C, a novel regulator of Dvl signaling abutting RNA-processing bodies, controls cilia orientation and leftward flow. Development 136, 3019–3030, doi:136/17/3019 [pii] 10.1242/dev.038174 (2009).

10. Piazzon, N., Maisonneuve, C., Guilleret, I., Rotman, S. & Constam, D. B. Bicc1 links the regulation of cAMP signaling in polycystic kidneys to microRNA-induced gene silencing. J Mol Cell Biol 4, 398–408, doi:10.1093/jmcb/mjs027 (2012).

11. Rothe, B. et al. Bicc1 Polymerization Regulates the Localization and Silencing of Bound mRNA. Mol Cell Biol 35, 3339–3353, doi:10.1128/MCB.00341-15 (2015).

12. Doidge, R., Mittal, S., Aslam, A. & Winkler, G. S. Deadenylation of cytoplasmic mRNA by the mammalian Ccr4-Not complex. Biochem Soc Trans 40, 896–901, doi:10.1042/BST20120074 (2012).

13. Collart, M. A. & Panasenko, O. O. The Ccr4--not complex. Gene 492, 42–53, doi:10.1016/j.gene.2011.09.033 (2012).

14. Chicoine, J. et al. Bicaudal-C recruits CCR4-NOT deadenylase to target mRNAs and regulates oogenesis, cytoskeletal organization, and its own expression. Dev Cell 13, 691–704, doi:10.1016/j.devcel.2007.10.002 (2007).

15. Suzuki, T. et al. Postnatal liver functional maturation requires Cnot complex-mediated decay of mRNAs encoding cell cycle and immature liver genes. Development 146, doi:10.1242/dev.168146 (2019).

16. Morita, M. et al. Depletion of mammalian CCR4b deadenylase triggers elevation of the p27Kip1 mRNA level and impairs cell growth. Mol Cell Biol 27, 4980–4990, doi:10.1128/MCB.02304-06 (2007).

17. Shinohara, K. et al. Two rotating cilia in the node cavity are sufficient to break left-right symmetry in the mouse embryo. Nat Commun 3, 622, doi:10.1038/ncomms1624 (2012).

18. Supp, D. M., Witte, D. P., Potter, S. S. & Brueckner, M. Mutation of an axonemal dynein affects left-right asymmetry in inversus viscerum mice. Nature 389, 963–966. (1997).

19. Nonaka, S., Shiratori, H., Saijoh, Y. & Hamada, H. Determination of left-right patterning of the mouse embryo by artificial nodal flow. Nature 418, 96–99 (2002).

20. Ma, M., Gallagher, A. R. & Somlo, S. Ciliary Mechanisms of Cyst Formation in Polycystic Kidney Disease. Cold Spring Harb Perspect Biol 9, doi:10.1101/cshperspect.a028209 (2017).

21. Thastrup, O., Cullen, P. J., Drobak, B. K., Hanley, M. R. & Dawson, A. P. Thapsigargin, a tumor promoter, discharges intracellular Ca2+ stores by specific inhibition of the endoplasmic reticulum Ca2(+)-ATPase. Proc Natl Acad Sci U S A 87, 2466–2470, doi:10.1073/pnas.87.7.2466 (1990).

22. Nicastro, G., Taylor, I. A. & Ramos, A. KH-RNA interactions: back in the groove. Curr Opin Struct Biol 30, 63–70, doi:10.1016/j.sbi.2015.01.002 (2015).

23. Nakamura, T. et al. Fluid flow and interlinked feedback loops establish left-right asymmetric decay of Cerl2 mRNA. Nat Commun 3, 1322, doi:10.1038/ncomms2319 (2012).

24. Hollingworth, D. et al. KH domains with impaired nucleic acid binding as a tool for functional analysis. Nucleic Acids Res 40, 6873–6886, doi:10.1093/nar/gks368 (2012).

25. Dowdle, M. E. et al. A single KH domain in Bicaudal-C links mRNA binding and translational repression functions to maternal development. Development 146, doi:10.1242/dev.172486 (2019).

26. Lambert, N. et al. RNA Bind-n-Seq: quantitative assessment of the sequence and structural binding specificity of RNA binding proteins. Mol Cell 54, 887–900, doi:10.1016/j.molcel.2014.04.016 (2014).

27. Zhang, Y. et al. Bicaudal-C spatially controls translation of vertebrate maternal mRNAs. RNA 19, 1575–1582, doi:10.1261/rna.041665.113 (2013).

28. Leal-Esteban, L. C., Rothe, B., Fortier, S., Isenschmid, M. & Constam, D. B. Role of Bicaudal C1 in renal gluconeogenesis and its novel interaction with the CTLH complex. PLoS Genet 14, e1007487, doi:10.1371/journal.pgen.1007487 (2018).

29. Li, X. et al. Adipocyte-specific disruption of mouse Cnot3 causes lipodystrophy. FEBS Lett 591, 358–368, doi:10.1002/1873-3468.12550 (2017).

30. Ukita, K. et al. Wnt signaling maintains the notochord fate for progenitor cells and supports the posterior extension of the notochord. Mech Dev 126, 791–803, doi:10.1016/j.mod.2009.08.003 (2009).

31. Zhang, Y., Park, S., Blaser, S. & Sheets, M. D. Determinants of RNA binding and translational repression by the Bicaudal-C regulatory protein. J Biol Chem 289, 7497–7504, doi:10.1074/jbc.M113.526426 (2014).

32. Park, S., Blaser, S., Marchal, M. A., Houston, D. W. & Sheets, M. D. A gradient of maternal Bicaudal-C controls vertebrate embryogenesis via translational repression of mRNAs encoding cell fate regulators. Development 143, 864–871, doi:10.1242/dev.131359 (2016).

33. Tran, U. et al. The RNA-binding protein bicaudal C regulates polycystin 2 in the kidney by antagonizing miR-17 activity. Development 137, 1107–1116, doi:137/7/1107 [pii] 10.1242/dev.046045 (2010).

34. Iwakawa, H. O. & Tomari, Y. The Functions of MicroRNAs: mRNA Decay and Translational Repression. Trends Cell Biol 25, 651–665, doi:10.1016/j.tcb.2015.07.011 (2015).

35. Behm-Ansmant, I. et al. mRNA degradation by miRNAs and GW182 requires both CCR4:NOT deadenylase and DCP1:DCP2 decapping complexes. Genes Dev 20, 1885–1898, doi:10.1101/gad.1424106 (2006).

36. Jonas, S. & Izaurralde, E. Towards a molecular understanding of microRNA-mediated gene silencing. Nat Rev Genet 16, 421–433, doi:10.1038/nrg3965 (2015).

37. Takao, D. et al. Asymmetric distribution of dynamic calcium signals in the node of mouse embryo during left-right axis formation. Dev Biol 376, 23–30, doi:S0012-1606(13)00036-5 [pii] 10.1016/j.ydbio.2013.01.018 (2013).

38. Mochizuki, T. et al. Cloning of inv, a gene that controls left/right asymmetry and kidney development. Nature 395, 177–181 (1998).

39. Watanabe, D. et al. The left-right determinant Inversin is a component of node monocilia and other 9+0 cilia. Development 130, 1725–1734 (2003).

40. Wilkinson, D. G. & Nieto, M. A. Detection of messenger RNA by in situ hybridization to tissue sections and whole mounts. Methods Enzymol 225, 361–373 (1993).

41. Minegishi, K. et al. A Wnt5 Activity Asymmetry and Intercellular Signaling via PCP Proteins Polarize Node Cells for Left-Right Symmetry Breaking. Dev Cell 40, 439–452 e434, doi:10.1016/j.devcel.2017.02.010 (2017).

42. Hashimoto, M. et al. Planar polarization of node cells determines the rotational axis of node cilia. Nat Cell Biol 12, 170–176, doi:ncb2020 [pii] 10.1038/ncb2020 (2010).

