## SUPPLEMENTAL INFORMATION for "Fluid flow-induced left-right asymmetric decay of *Dand5* mRNA in the mouse embryo requires Bicc1-Ccr4 RNA degradation complex"

Supplementary S1 Information for Fig3

**a**

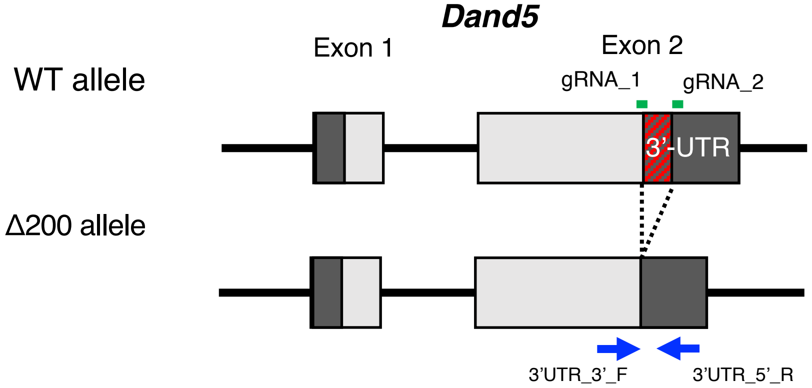


**b**

### Exon 2 of *Dand5*

gtgatctccaggcctggttgcacaagtgcccgggtccttaatcatctctgttttggccgctgttcctccttctacattcccagctcg
gatcccacccctgtagtcttctgcaacagctgtgtgccggctcgaaagcgctggacatcggtgacgctgtggtgtggagctg
gccaattagcctcccctcggcgggtgaggatttccacggtattggtccagaagtgtcagtgccgc*ccgaagctgtgaGCT*
*GAGCATCCTAGAGG*AATGCGCAGGACATGAATGAACCTTGGCAAGAAGCAGGAGAC
GCAATAGAAAGACGTGACCTGAATGATGTGCATCGGGTCAAGAGAGCAGAATTGGA
CAGGGCCAGAATACTAGACATGGTCCCCCAGTCATGGGTTTAGACCACTAATAGTCG
TAGAATGTGGGGCTAGGGTATAACTCAGTGGGAGAGGGCTTGCCTAGCATGCAtgaag
*ccctgggttctattcgtatgtga*tatgtggaggttaaaaaaaggtgaaaattagctatagtgtatgattagccctgtgtgagagg
gaacattcatgccacaatcactaactcacgtcctcctcaggaatcctcagcagtcccgcaaggtagaatttcaccatgggtc
attttacagatgaggaaaccgaagctatttaggcactgtcacttttggagaggggtggaatagggagagggcttcagacac
agccctgacgcacgcgcagcacagctccctgctgagctcctcccccagcctggggttccgcttttagccactaggtggcac
cgcgtcccagtgtccaagacagacactcccaactgttatgatgacagcactacccaaacctggacactcagcaaagatga
ccctgttctggggacaccagaataacagcagcagcagcaacaacaacagcaacaaactatggcaggcaagcaggaa
gcagctgctaggaaaacctggccccaaccagaggcgaccccgcatgtgtgcagctgcctctctcatcgacagtagccag
gggtttccctttttgccccacccgagaccccgcagcagcctcatgcccagcctggcattccgtgggaaccataaaggttggc
caagtcc

Target sequences
gRNA_1: ctgtgaGCTGAGCATCCTAGAGG
gRNA_2: tcacatacgaatagaacccaggg

3'-UTR
Red : Target sequence (gRNA) + PAM sequence
Expected deletion sequence

**c**

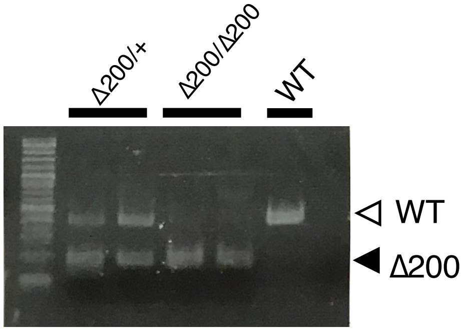


PCR primers
3'UTR_3'_F: GTGTGGAGCTGGCCAATTA
3'UTR_5'_R: CTCACACAGGGCTAATCATACA

FigS2 elated to Fig4

**a**

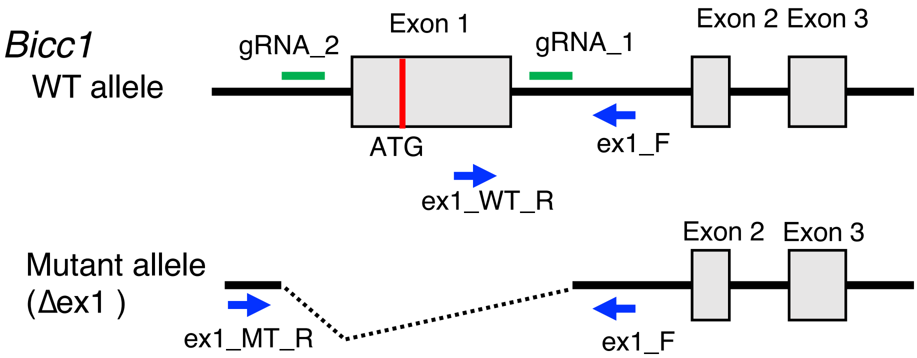


Target sequences

gRNA_1: GAGGGAGGGTGTCCCGTGAGCGG

gRNA_2: AGCCCTGCGCATCCCCGCCTCGG

PCR primers

ex1_F: 5'-TAAGGAGGGGAGGGAAAAGA

ex1_WT_R: 5'-GCGTGGACAGGAAGAAACTC

ex1_MT_R: 5'-CTAGGCTCGAACTGGGTGAG

Modified sequence of *Bicc1* mutant line

Line #2: CCCGTGGGCCTGGCCGGCCAATGAGCG TCCCT CCCCCCCAACTTTGTCAGCTTGCTC

**b**

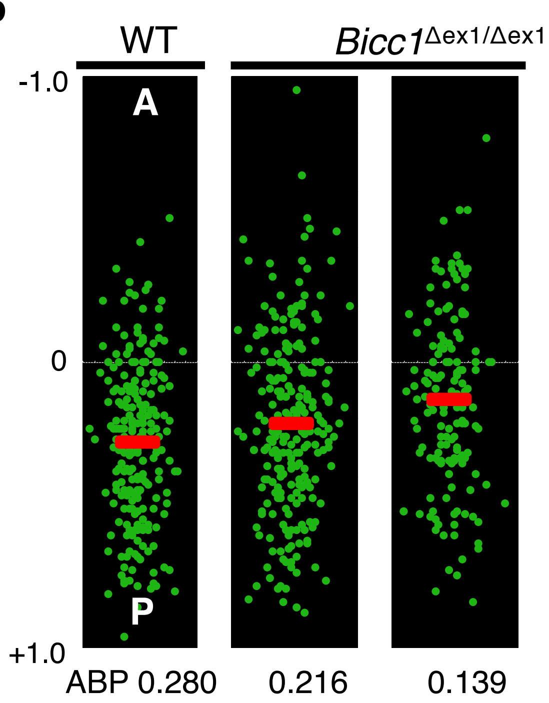


**c**

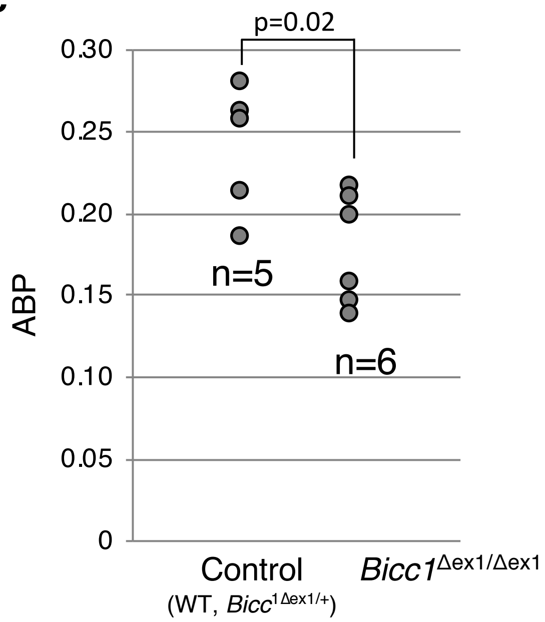

FigS3 related to Fig5

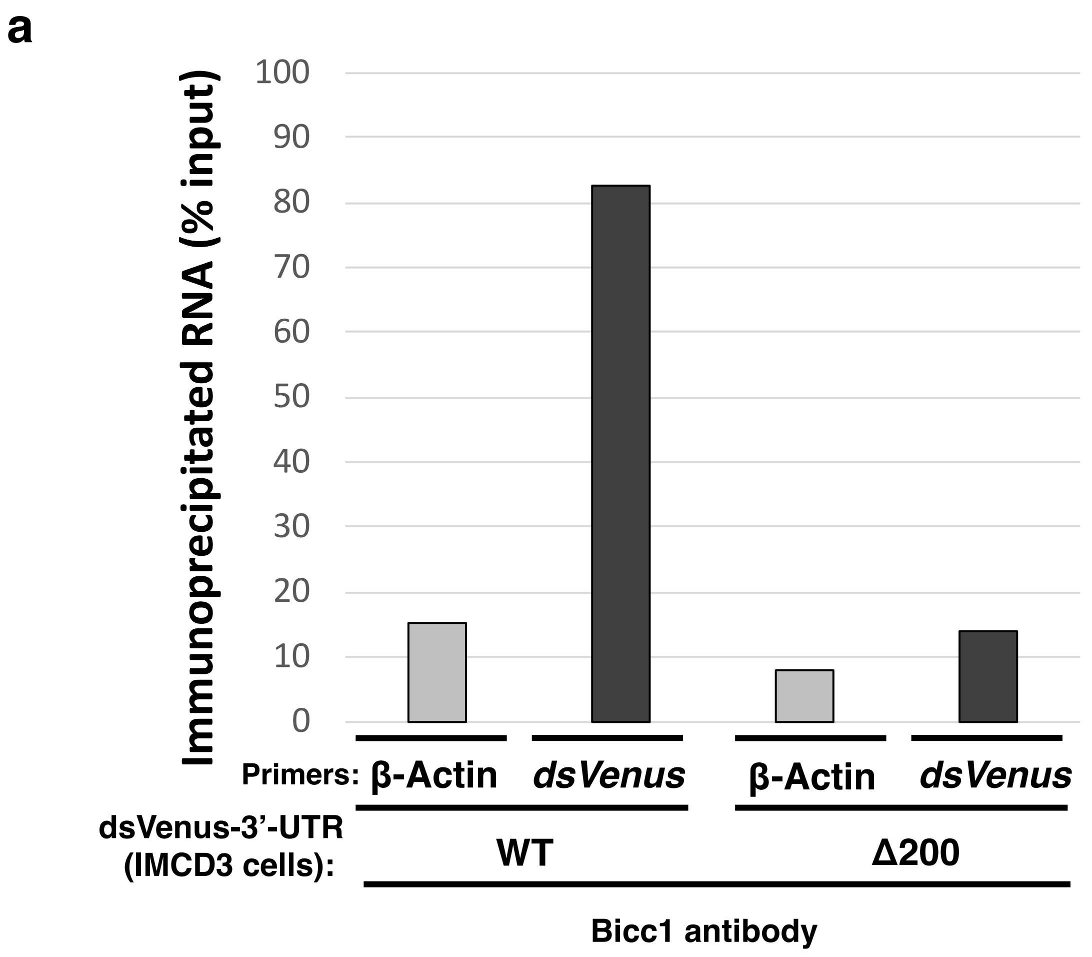


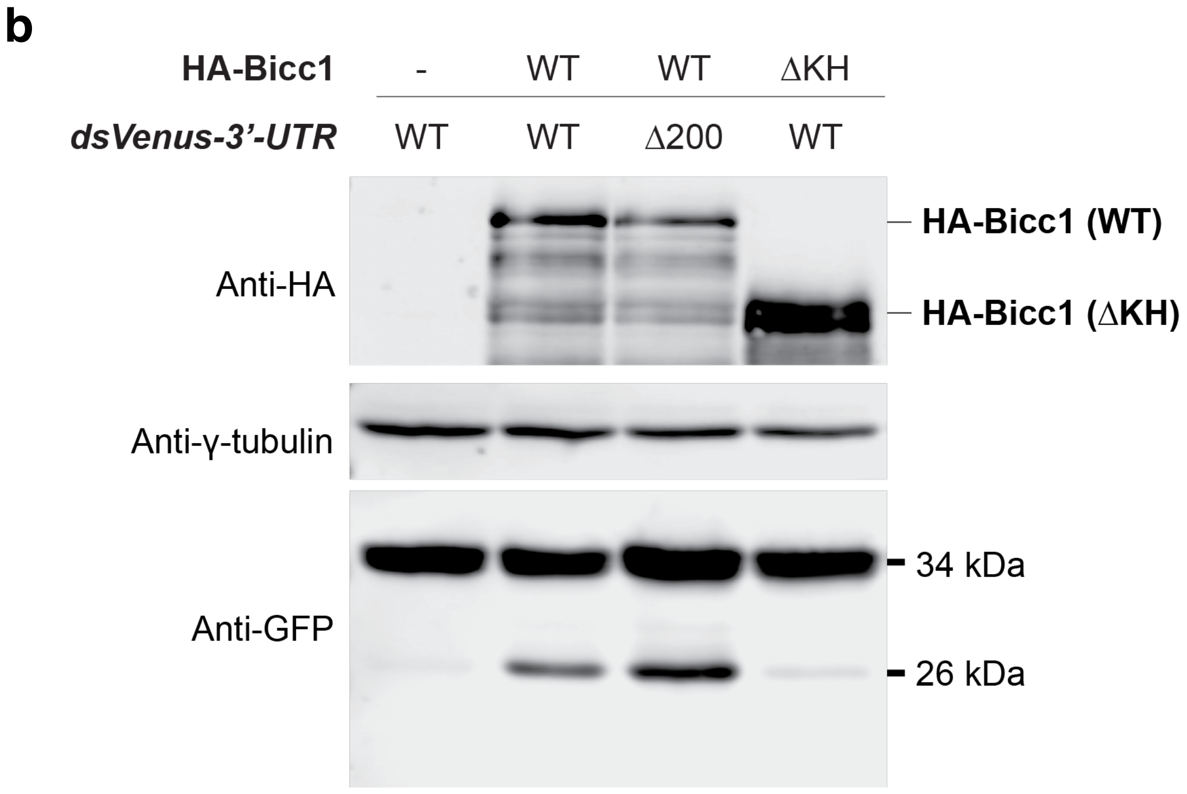

**SUPPLEMENTAL INFORMATION**

**Supplementary Figure 1. Generation of Mice with the *Dand5*$^{\Delta 200}$ Allele with the CRISPR/Cas9 System**

**a** Schematic representation of deletion of the 200-bp DNA sequence corresponding to the proximal-most region of the 3'-UTR of *Dand5* mRNA from the mouse genome. The red-shaded region of the WT allele was deleted to give rise to the *Dand5*$^{\Delta 200}$ allele. The positions of guide RNAs (gRNAs, green bars) and of PCR primers for genotyping (blue arrows) are indicated. **b** Nucleotide sequence of exon 2 of mouse *Dand5* showing portions related to deletion of the 200-bp fragment in (A) with the CRISPR/Cas9 system. **c** PCR-based genotyping of WT, *Dand5*$^{\Delta 200/+}$, and *Dand5*$^{\Delta 200/\Delta 200}$ mouse embryos with the indicated primers.

**Supplementary Figure 2. Generation of *Bicc1* Mutant Mice with the CRISPR/Cas9 System**

**a** *Bicc1* mutant mice were generated by CRISPR/Cas9 editing with two gRNAs (green bars) to delete exon 1. The positions of PCR primers (blue arrows) for genotyping are also indicated. **b** Relative position of the basal body in node cells of WT and *Bicc1*$^{\Delta ex1/\Delta ex1}$ embryos at three- to five-somite stages. Each green dot corresponds to the relative position of one basal body along the anterior (A)–posterior (P) axis. The average basal body position (ABP) for each embryo is indicated by the red bar, with the corresponding value being given below each image. **c** Summary of ABP for the indicated numbers of embryos (n) determined as in (B). The p value was determined with the T test.

**Supplementary Figure 3. Binding of Endogenous Bicc1 to the *dsVenus-3'-UTR* Reporter mRNA**

**a** A cytoplasmic fraction prepared from IMCD3 cells stably expressing the *dsVenus-3'-UTR* or *dsVenus-3'-UTRΔ200* constructs was subjected to immunoprecipitation with antibodies to Bicc1, and the resulting immunoprecipitates as well as a portion of the input material were subjected to RT-qPCR analysis with PCR primers specific for dsVenus or β-actin (negative control). Data represent the ratio of co-immunoprecipitated mRNA to the input. **b** Binding of Bicc1 to the *dsVenus-3'-UTR* Reporter mRNA Does Not Affect the Abundance of the Encoded Protein. Cytoplasmic extracts of HEK293T cells transfected as in Figure 5B were subjected to immunoblot analysis with antibodies to GFP (for detection of dsVenus), to HA (for detection of HA-

Bicc1), and to γ-tubulin (loading control). The antibodies to GFP detected both the intact dsVenus protein (34 kDa) and a degradation product (26 kDa).
